# Global mapping of VEGF-A driven changes in the RNA bound phospho-proteome reveals a molecular switch to coordinate angiogenesis in endothelial cells

**DOI:** 10.64898/2026.11.11.765910

**Authors:** Rhys Wardman, Anna Tikhonova, Katrin Bennewitz, Lukas Hermann, Shubhang Joshi, Jennifer Schwartz, Frank Stein, Roxana Ola, Gergana Dobreva, Annette Hammes, Jens Kroll, Joerg Heineke

## Abstract

RNA binding proteins (RBPs) serve as upstream master regulators of cellular fate transitions at the post-transcriptional level. RBP-RNA interactions are dynamically regulated by post-translational modifications to coordinate cellular responses. We hypothesised that changes in RBP phosphorylation will play a key role in coordinating angiogenesis, the formation of new blood vessels, in endothelial cells (ECs). Therefore, we used phospho-proteomic analysis of the mRNA bound proteome to identify global phosphorylation changes in mouse cardiac ECs during acute and prolonged VEGF-A stimulation. Of the differentially phosphorylated RBPs, we found that the phosphorylation of serine 199 within mRNA bound Phf6 (PHD finger protein 6) was consistently increased following VEGF-A stimulation. Phf6 phosphorylation at this specific site was sufficient and required to induce a pro-angiogenic phenotype, facilitating a change in Phf6 conformation promoting interaction with distinct RNAs. We show here that Phf6 phosphorylation at serine 199, downstream of VEGF-A, acts as a molecular switch to differentially coordinate a decrease anti-angiogenic RNAs (i.e. extracellular matrix genes) and an increase in pro-angiogenesis related RNAs, in part by differentially regulating their stability, to promote angiogenesis.

**GRAPHICAL ABSTRACT:** 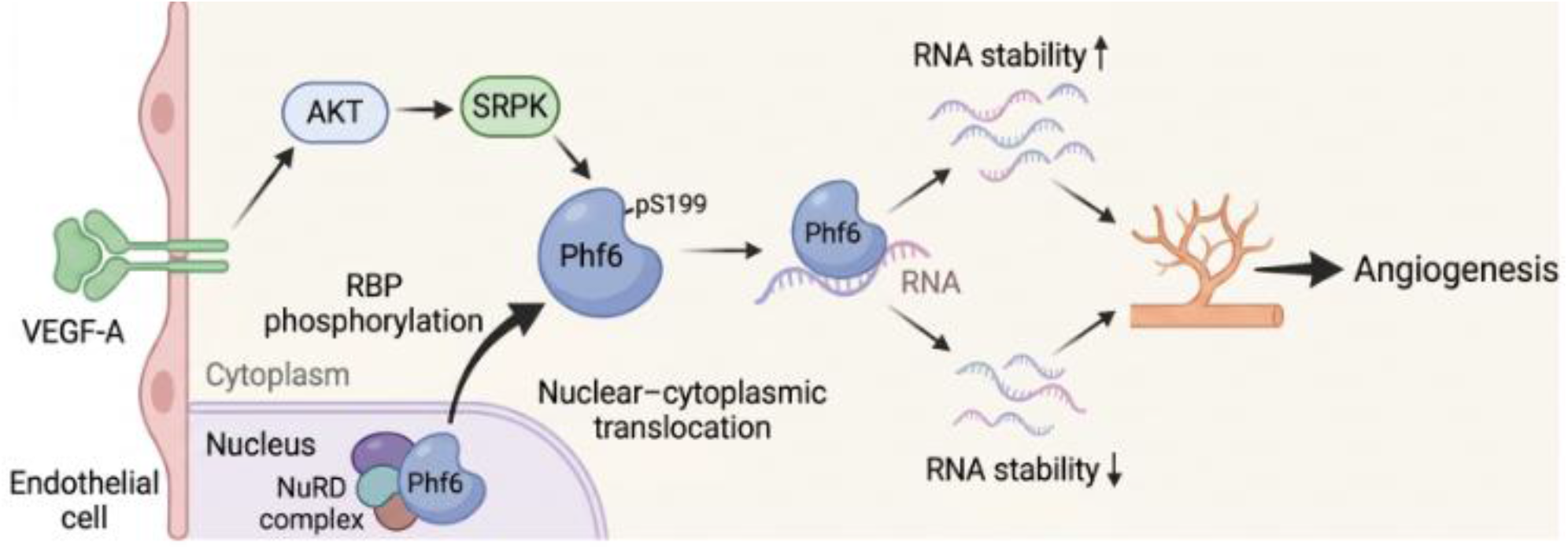

## INTRODUCTION

The interaction between RNA binding proteins (RBPs) and their target subsets of RNAs are highly dynamic, allowing distinct changes in RNA-protein interactions to serve as a primary response mechanism to alterations in cellular conditions. Variations in RNA-protein interactions are determined by a range of factors such as changes in the abundance of the target RNAs, post-translational modifications in the RBP, changes in RBP (or target RNA) localisation, modifications to subsets of target RNAs such as N6-methyladenosine (m⁶A) or A-to-I editing, or binding competition with other proteins (1,2,3,4). Of these modifications, post-translational modifications to the RBP (such as phosphorylation, methylation or acetylation) are perhaps the most classical drivers of changes in RBP function in response to specific signalling cues (2,5).

Phosphorylation of an RBP can couple extracellular signalling prompts, via downstream kinase activation, to coordinated and specific changes in binding affinity for subsets of target RNAs. This is achieved, for example, by prompting structural changes (i.e. change in conformation or charge) within the protein, changes in localisation, protein-protein interactions, or stability, and has been implicated in numerous physiological and pathophysiological responses (6,7,8). This altered RBP activity serves to differentially sculpt the cellular proteome such as in altering the translation, splicing, localisation or stability of target RNAs.

Endothelial cells (ECs) form the border between circulating blood and underlying tissues, playing a crucial role in nutrient and oxygen delivery. ECs serve as front line responders to physiological and pathophysiological changes in conditions, such as increased growth factor abundance, changes in circulating metabolites or oxygen tensions as well as emerging inflammatory mediators and processes (9,10). Given their highly dynamic and heterogeneous nature, ECs must remain primed to quickly respond to external cues. We propose that changes in RNA-protein interactions play a key role in facilitating this EC plasticity at the post-transcriptional level. For example, in response to transforming growth factor-β stimulation, RBPs including heterogeneous nuclear ribonucleoprotein H1 (hnRNP H1) and cold shock domain containing E1 (Csde1) actively maintain EC function and counteract mesenchymal activation (11). Others have also shown key roles of specific RBPs in regulating EC biology, for example the RBP Quaking (QKI) regulates EC cell cycle progression and maintain EC barrier function, while HuR (Human antigen R/ELAVL1) is an RBP shown to coordinate mechanical and biochemical stress responses in ECs (12,13).

Angiogenesis (the formation of new vasculature from pre-existing vessels) is an EC led process promoted primarily by vascular endothelial growth factor A (VEGF-A), which drives a number of phenotypic alterations such as enhanced EC proliferation and migration to support neo-vascularisation (14,15,16). Angiogenesis is differentially regulated under physiological or pathological conditions with both beneficial or detrimental effects. For example, selective stimulation of angiogenesis in the heart is a promising therapeutic avenue to improve repair following myocardial infarction, whereas exaggerated, abnormal angiogenesis is a hallmark of cancer, facilitating tumour growth and metastasis (16,17,18,19, 20). Therefore, there is a need to understand the specific regulators of the angiogenic program in ECs in order to more effectively target the angiogenic process therapeutically.

Recent studies have demonstrated the global effects of RBP phosphorylation on RNA occupancy, showing that widespread regulatory RBP phosphorylation networks can act to coordinate RBP function (8). Here, we set out to build on this by integrating RNA interactome capture (oligo d(T) precipitation of UV-crosslinked RNA–protein complexes) with phospho-proteomic analyses to uncover, for the first time, how growth factor stimulation (VEGF-A) drives global changes in RBP phosphorylation to orchestrate cellular signalling responses.

Using the model of VEGF-A induced angiogenesis in mouse cardiac endothelial cells (MCECs), we identified global VEGF-A driven changes in the phosphorylation of the mRNA bound proteome and characterised the functional consequences of specific differential phosphorylation events. We found that the phosphorylation of PHD Finger Protein 6 (Phf6) at Ser199 downstream of VEGF-A drives a specific interaction between Phf6 and angiogenic or mesenchymal RNAs sufficient and required to coordinate a pro-angiogenic phenotype.

In its canonical function, Phf6 is a chromatin-associated transcriptional regulator that interacts with chromatin remodelling and transcriptional complexes and has essential roles in development, haematopoiesis, and tumour suppression (21,22,23,24). Recently, Phf6 has emerged as an mRNA binding protein, which was identified in multiple global RNA interactome capture screens (25,26,27,28). Here, we show that VEGF-A driven Phf6 Ser199 phosphorylation induces specific changes in the localisation and RNA binding of Phf6 to distinct oligonucleotide motifs and RNA targets involved in the regulation of angiogenesis in murine and human ECs. This results in a switch from the canonical function of Phf6 as a transcriptional regulator in the nucleus to a cytoplasmic RBP, where it acts to differentially orchestrate a proangiogenic phenotype by decreasing the expression of anti-angiogenic RNAs by binding to their 3’ UTR and decreasing their stability, whilst increasing the expression of proangiogenic RNAs.

## MATERIAL AND METHODS

### Cell models and culture

All cells were cultured under standard conditions (5% CO_2_, 37⁰C).

Mouse cardiac endothelial cells (MCECs, CELLutions biosystems inc. Cat# CLU510) were used between P5-P20, and were cultured in high glucose (4.5 g/L) DMEM supplemented with 5 % foetal calf serum (FCS).

Human umbilical vein endothelial cells (HUVECs) were obtained from PromoCell (Cat# C-12200) and cultured in PromoCell EC growth media with growth media supplement mix (Cat# C-22110). HUVECs were used between P2-6.

AKT overexpression in HUVECs was achieved by lentiviral transduction as published in 29. hiPSC-derived ECs were obtained as previously published (30). Briefly, 6 cm dishes were precoated with Matrigel (Corning, Cat#354230) (1:100 dilution in cold PBS) and incubated at 37°C overnight. hiPSCs (Allen Institute for Cell Science, AICS-0052-003, WTC-mEGFP-MYL7-cl3, tagged with CDH5-tomato, 31), were seeded on Matrigel-coated dishes in Essential 8 media (Thermo Fisher Scientific, Cat#A1517001) with 5 μM Y27632 (Miltenyi Biotec, Cat# 130-106-538). The medium was changed to Essential 8 without 5 μM Y27632 after 16-24 hours and changed every day until they achieved 75–85% confluency. Thereafter, the cells were washed twice with PBS and incubated at 37°C for 1 minute in PBS-EDTA buffer (500 ml PBS, 500 µl 0.5 M EDTA, 0.9 g NaCl). Then, PBS-EDTA buffer was aspirated, and Essential 8 media + 1 µM CHIR-99021 (Selleckchem, Cat#S1263) + 5 µM Y27632 was added. The cells were scraped and ∼1.6 × 10^5^ cells cm^2^ were seeded in a Matrigel-coated 24-well plate. To start the differentiation, the media was changed to RPMI (Thermo Fisher Scientific, Cat#61870010) + 100 ng ml−1 activin A (R&D Systems, Cas#338-AC) + 1× B-27 minus insulin (Thermo Fisher Scientific, Cas#A1895601) + Matrigel (1:60 dilution).

After 17h of incubation at 37°C, the media was changed to RPMI + 5 ng ml−1 BMP4 (R&D Systems, Cas#314-BP) + 1 µM CHIR-99021 + 1× B-27 minus insulin to generate cardiogenic mesoderm cells. After 24 hours, the media was changed to StemPro (Thermo Fisher Scientific, Cas#10639011) + 4 × 10−4 M MTG + 2 mM l-glutamine (Thermo Fisher Scientific, Cat#25030024) + 50 ug ml−1 ascorbic acid (Merck, Cat#A4544) + 10 ng ml−1 BMP4 + 5 ng ml−1 bFGF (R&D Systems, Cat#233-FB) + 300 ng ml−1 VEGF (R&D Systems, Cat#293-VE-500) and the cells were incubated for 72 hours to generate cardiac mesoderm-derived ECs. The cells were split on day 5 and reseeded in EGM (Lonza, Cat#CC-3124) + 20 ng ml−1 VEGF + 20 ng ml−1 bFGF + 1 µM CHIR-99021 at a density of 2 × 104 cells cm−2 on 0.1% (vol/vol) gelatin-coated plates. Thereafter, the cells were split when they achieved 75-85% confluency until day 14 to generate definitive cardiac-ECs. Cells were washed in PBS and incubated in serum free media overnight before stimulation. Stimulation conditions are indicated in the respective Figure legends.

### RNA interactome capture

RNA interactome capture was performed as we have previously published (11) and as detailed in 32, with only minor changes. Briefly, cells (four 10 cm plates per replicate) were cultured until ∼80% confluent and before stimulation under the respective conditions. Cells were washed in chilled PBS, which was fully aspirated and protein-RNA complexes cross-linked by 254 nm UV radiation (150 mJ/cm^2^). Cells were lysed (500 mM LiCl, 0.5% w/v LiDS, 5 mM DTT, 1 mM EDTA 20 mM, cOmplete™ EDTA-free Protease Inhibitor Cocktail (Roche) and Phosphatase Inhibitor Cocktail (Merck, Cat# 524629)) and homogenised with a 21-gauge needle. Protein-RNA complexes were captured using Oligo d(T)25 magnetic beads (NEB Cat# S1419S) for 2 hours at 4⁰C followed by magnetic separation. Beads were washed in lysis buffer, twice in wash buffer I (500 mM LiCl, 0.1% LiDS, 5 mM DTT, 1 mM EDTA, 20 mM Tris-HCl pH 7.4), twice in wash buffer II (500 mM LiCl, 5 mM DTT, 1 mM EDTA, 20 mM Tris-HCl pH 7.4) and twice in low salt buffer (200 mM LiCl, 5 mM DTT, 1 mM EDTA, 20 mM Tris-HCl pH 7.4). Residual buffer was removed by centrifugation and RNA-protein complexes were eluted (75°C for 5 minutes) in elution buffer. Proteins were released by RNase A/T1 digestion (1 h at 37°C).

### Proteomic/Phosphoproteomic analysis

Proteomic and phosphoproteomic analysis was performed as in 33. RNA binding proteins were isolated by RNA interactome capture as above. For analysis, each sample was divided in two for simultaneous identification. Proteins were processed via SP3 on a KingFisher Apex™ system (Thermo Fisher). Digestion used trypsin (1:20 protease:protein) in 50 mM HEPES, 5 mM TCEP, 20 mM CAA for 5 h at 37°C. Up to 10 µg peptides were labelled with TMT10plex™: 4 µL of TMT reagent (0.8 mg in 45 µL acetonitrile) was added per sample, incubated 1 h at room temperature, and quenched with 4 µL 5% hydroxylamine (15 min, RT). Labelled samples were multiplexed, desalted (Oasis® HLB µElution Plate, Waters), and vacuum-dried. Phosphopeptide enrichment followed 34: peptides were resuspended in IMAC loading solvent (80% acetonitrile, 0.4% TFA) and enriched on the KingFisher Apex™ using Fe-NTA magnetic beads (Cube Biotech), eluting with 0.2% dimethylamine in 80% acetonitrile before drying (34). The enrichment supernatant was fractionated offline by high-pH reversed-phase HPLC (Agilent 1200 Infinity, Gemini C18 column, 3 µm/110 Å, 100 × 1.0 mm, with Gemini C18 SecurityGuard pre-column, Phenomenex). Mobile phases were 20 mM ammonium formate, pH 10.0 (A) and 100% acetonitrile (B); at 0.1 mL/min: 100% A for 2 min, to 35% B over 59 min, to 85% B over 1 min, hold 15 min, re-equilibrate at 100% A for 13 min. 48 fractions were collected and pooled into 6.

Samples were analysed on an UltiMate 3000 RSLCnano (Thermo Fisher) with a C18 PepMap™ 100 trapping cartridge (300 µm × 5 mm, 5 µm, 100 Å) and a nanoEase™ M/Z HSS T3 analytical column (75 µm × 250 mm, 1.8 µm, 100 Å, Waters). Peptides were trapped at 30 µL/min in 0.05% TFA (6 min), then eluted at 0.3 µL/min with increasing solvent B (3% DMSO, 0.1% FA in acetonitrile) against solvent A (3% DMSO, 0.1% FA in water). Peptides were sprayed (2.4 kV, Pico-Tip emitter, 360 µm OD × 20 µm ID, 10 µm tip, CoAnn Technologies) into an Orbitrap Fusion™ Lumos™ Tribrid™ (Thermo Fisher) at 275°C capillary temperature.

Full MS scans: m/z 375–1,500 (to 1,650 for phospho samples), 120,000 resolution at m/z 200, 50 ms max injection, ’standard’ AGC. DDA mode with MS/MS in the Orbitrap at 30,000 resolution, 94 ms (110 ms) max injection, 200% AGC target, HCD fragmentation at 36% NCE, profile-mode MS2. Quadrupole isolation window: 0.7 m/z; dynamic exclusion: 60 s (20 s); precursor charge states 2–7 only. Raw files were converted to mzML (MSConvert, ProteoWizard; peak picking, 64-bit encoding, zlib compression, top 1000 peaks) and searched with MSFragger v3.8 in FragPipe v20.0 against UP000000589 (Mus musculus, ID 10090, 21,928 entries, October 2022) with contaminants and reversed sequences. Fixed modifications: carbamidomethylation (C, +57.0215), TMT10 (K, +229.1629). Variable modifications: oxidation (M, +15.9949), N-terminal acetylation (+42.0106), N-terminal TMT10 (+229.1629); phospho searches additionally included pSTY (+79.9663). Mass tolerance: 20 ppm (MS1 and MS2). Trypsin, ≤2 missed cleavages, minimum peptide length 7. FDR threshold: 0.01 (peptide and protein level).

Peptide/PTM-level (psm.tsv) and protein-level (protein.tsv) data from FragPipe were processed in R, excluding contaminants and reverse hits (35). Phosphopeptides were filtered by STY.79.9663.Best.Localization ≥ 0; proteins required ≥2 razor peptides (Razor.Peptides ≥ 2). Each phosphopeptide was assigned a unique identifier (gene name + modified sequence, with asterisks marking modified residues, followed by site count and positions), and PSMs sharing an identifier within a sample were summed to yield one TMT intensity per peptide per sample (3,718 phosphopeptides passed QC). Phosphopeptide intensities were normalised to protein abundance: for each gene/condition, the phosphopeptide:protein intensity ratio was corrected by the gene-specific median ratio, then rescaled to the median phosphopeptide intensity to preserve interpretable units (’nonmod.norm.phospho’). Log2-transformed values were assembled into a Bioconductor SummarizedExperiment. Batch effects were removed from log2 raw TMT intensities using limma’s removeBatchEffect (36), followed by VSN normalization (normalizeVSN) (37). For visualization, abundances were scaled to the median of control replicates. Differential expression used limma’s moderated t-test, with replicate included as a design factor; where limma yielded no significant hits, fdrtool (38) was applied to the t-values as a more sensitive alternative. Peptides were called hits at FDR < 0.05 and |fold change| > 2, and candidates at FDR < 0.2 and |fold change| > 1.5. Hits and candidates were clustered by k-means (Euclidean distance, ward.D2 linkage) on control-normalized median abundances.

### Transfections

For protein/construct overexpression, cells were transfected with Lipofectamine 3000 (Invitrogen) as per the manufacturer’s instructions. Constructs were transfected at a final concentration of 2 µg/ml.

Flag-SRm160 was a gift from Phil Sharp (Addgene plasmid # 11305, 39). Myc-Phf6 was purchased from Origene (MR205605) and transformed into DH5α competent cells.

siRNA transfection was performed with Lipofectamine RNAiMAX according to the manufacturer’s instructions using 20 nM (MCECs) or 40 nM (HUVECs) siRNA for 48 hours. The following siRNAs were used in the manuscript; Phf6 mouse (ON-TARGETplus Mouse Phf6 siRNA Dharmacon L-058690-01), Phf6 human (ON-TARGETplus Human PHF6 siRNA Dharmacon L-014862-00), Srrm1 mouse (ON-TARGETplus Mouse Srrm1 siRNA Dharmacon L-045815), Notch2 human (ON-TARGETplus Human NOTCH2 siRNA Dharmacon L-012235-00), Col5a1 human (ON-TARGETplus Human COL5A1 siRNA Dharmacon L-011015-00).

### Immunoprecipitation

Cells (two 10 cm plates per replicate) were lysed in lysis buffer (50 mM Tris HCl (pH 7.4), 100mM NaCl, 1% NP-40, 0.1% SDS, 0.5% sodium deoxycholate, cOmplete™ EDTA-free Protease Inhibitor Cocktail (Roche) and Phosphatase Inhibitor Cocktail (Merck).

For IP of endogenous protein (Phf6), 50 µl of Magne® Protein G beads (Promega) were coupled to 2 mg of anti-Phf6 antibody (Proteintech, Cat# 51062-1-AP) overnight at 4°C. For IP of Myc and Flag tagged constructs, 25 µl of magnetic Myc-Trap (Proteintech Cat# yt2ma) or agarose DYKDDDDK Fab-Trap (Proteintech Cat# ffa) were used.

Lysate was combined with beads and incubated at 4°C for 2 hours. Lysate was removed, and beads were washed in high salt buffer (50 mM Tris-HCl (pH 7.4), 1 M NaCl, 1 mM EDTA, 1% NP-40, 0.1% SDS, 0.5% sodium deoxycholate) for five minutes followed by three five-minute washes in wash buffer (20 mM Tris-HCl (pH 7.4), 10 mM MgCl2, 0.2% Tween 20). Samples were eluted in 20 µl of Laemmli Sample Buffer by heating at 95°C.

### Western blot

Western blots were carried out according to well established methodologies. Samples were heated in Laemmli sample buffer at 95°C for five minutes and separated on 10% polyacrylamide gels (using standard SDS-PAGE procedure). Proteins were transferred to PVDF membranes, and following blocking were incubated with corresponding primary antibodies overnight at 4°C.

The following primary antibodies were used: Phf6: Rabbit polyclonal antibody, Proteintech 51062-1-AP, 1:1000. Srrm1: Rabbit polyclonal antibody, Proteintech 12822-1-AP, 1:1000. Myc: Mouse monoclonal antibody, Cell Signalling, 2276S, 1:1000. Flag: Rabbit polyclonal antibody, Proteintech 20543-1-AP. Phosphoserine: Mouse monoclonal, Merk P5747 1:1000. hnRNP H1: Rabbit polyclonal antibody, Invitrogen PA5-50678, 1:1000. Erk 1/2: Rabbit polyclonal antibody Cell Signalling 9102, 1:1000. GAPDH: Mouse monoclonal antibody, Proteintech 60004, 1:1000. Lamin A/C Rabbit polyclonal antibody, Cell Signalling 2032, 1:1000.

After washing and incubation with the corresponding secondary antibodies (Anti-rabbit HRP Amersham NA934, 1:10,000 or Anti-mouse HRP Thermo Fisher 62-6520, 1:10,000), blots were imaged with an Amersham 680 imager. Images analysed with ImageStudio software (ver 5.2) using the analysis feature.

### Proximity ligation assay (PLA)

Cells were fixed in 4% paraformaldehyde and permeabilised in 1% BSA, 0.1% TritonX100 in PBS. For the phospho-serine PLA, cells were blocked in Duolink blocking buffer for 1 hour at 37°C. Primary antibodies (either Phf6; Proteintech 51062-1-AP, 1:500, or Srrm1; Proteintech 12822-1-AP, 1:500, both rabbit) were diluted in Duolink antibody dilutant and added to cells overnight at 4°C. Cells were washed in Duolink buffer A and incubated with Duolink anti-rabbit plus and anti-phosphoserine minus probes diluted 1:5 in antibody dilutant and incubated for 1 h at 37°C.

For the oligo d(T)/mRNA binding PLA, cells were washed in 0.1M triethanolamine and acetic anhydride added to 0.5%. Cells were washed in PBS 0.02% Tween20, and then in hybridisation buffer (50% deionised formamide, 5X SSC, 1 x Denhardts reagent, 0.1% (v/v) Tween20, 0.1% (w/v) CHAPs, 5 mM EDTA, 1 mg/ml tRNA and 100 µg/ml heparin).

Biotynylated oligo d(T) DNA probe ([BtnTg]TTTTTTTTTTTTTTTTNN) was diluted to 200 nM in hybridisation buffer, boiled at 95°C for five minutes and added to cells overnight at 37°C. Cells were washed in 50% deionized formamide/5×SSC, followed by 25% deionized formamide/1×SSC, 12.5% deionized formamide/2×SSC, 2×SSC/0.1% Tween20 and 0.2×SSC/0.1% Tween20. After a final wash in PBS 0.02% Tween20, cells were blocked in Duolink blocking buffer for 1 h at 37°C. Primary antibodies were diluted in Duolink antibody dilutent (Phf6 or Srrm1 rabbit antibody with mouse anti-biotin antibody,1:500 dilution) and added to cells overnight at 4°C. Cells were washed in Duolink buffer A and anti-Rabbit plus/anti-mouse minus probes diluted 1:5 in antibody dilutant and incubated for 1 h at 37°C.

Cells were washed in buffer A, and incubated with Duolink ligase mix for 30 mins followed by amplification mix (red) for 100 minutes hour at 37°C. Cells were washed in Duolink buffer B and mounted in Vectashield Mounting Medium with DAPI. For simultaneous staining of endogenous protein (Phf6 or Srrm1), and additional incubation step in anti-Rabbit Alexa-fluor 488 was included).

### Nuclear-cytoplasmic fractionation

Cells were washed twice in chilled PBS, scraped into 1 mL PBS and pelleted by centrifugation. This was followed by lysis in 1 mL 0.1% IGEPAL CA-630 in PBS (with collection of input samples). Remaining lysate was centrifuged and the cytoplasmic fraction was removed to a fresh tube. The nuclear pellet was washed in 0.1% IGEPAL CA-630 before lysis in Laemmli sample buffer.

### Tube formation assay

Cells (25000 cells per well) were plated onto 96 well plates precoated with Corning Matrigel basement membrane. Tube formation was assessed 24 hours later by addition of 1:5000 0.1% BCECF AM Ester (2’,7’-bis-(2-carboxyethyl)-5-(and-6)-carboxyfluorescein, acetoxymethyl ester) and fluorescent microscopy. The total number of complete loops per visible field were quantified, with each data point being the average of three images taken per well.

### EdU incorporation/proliferation assay

EdU labelling was perfomed using the Invitrogen Click-iT™ Plus EdU Cell Proliferation Kit as per the manufacturer’s instructions (Invitrogen C10637). Cells were plated onto glass coverslips and stimulated under the respective conditions. EdU was added to cells to a final concentration 10 µM for 2 hours at 37°C. Cells were then fixed in 4% paraformaldehyde for 15 minutes and permeabilised in 0.5% Triton X-100 for 20 minutes. Cells were then incubated in Click-iT™ reaction cocktail for 30 minutes at room temperature, as per the manufacturer’s instructions. Cells were washed, mounted in Vectashield Mounting Medium with DAPI, and visualised with fluorescent microscopy.

### Migration assay

Cells were plated onto a 6 well plate, premarked with a grid on the underside for orientation, and stimulated under the respective conditions. Mitomycin C (10 μg/mL) was added for 2 hours. Cells were scratched along the grid with a sterile tip, media was replaced and migration assessed by analysing closure after 24 hours using Image J software.

### Morpholino based knock-down and SRPK inhibition in zebrafish and hyaloid vasculature assessment

Morpholinos:

Two independent morpholinos were used to knock-down Phf6 expression; morpholino 1 (MO1) is a slice blocking morpholino sequence 5′TGTACAGCTAACATACCATGCACTT′3 (40) and MO2 is a translation blocking morpholino sequence 5′TGTCCTGACATGACCAGTTTTCATÁ3. Both were supplied by, and MO2 designed by, GeneTools, LLC.

The Morpholino stock solution (16 µg/µl) was heated at 65 °C for five minutes and diluted to 6 µg/µl with 0.1 M KCl for use. Freshly laid zebrafish eggs (Tg(fli1:EGFP) zebrafish) larvae were collected and arranged in a 1% agarose ramp in E3 medium. One nl of morpholino was then injected into the yolk sac of one-cell stage embryos. The larvae were collected for the hyaloid experiment at 120 hpf.

SRPK inhibition:

For SRPK inhibition, fertilized eggs were incubated at 28.5 °C in 6-well plate, containing around 30 embryos per well with 5 ml eggwater. This solution contained 0.003% PTU and either 500nM or 100nM of MSC-1186 (SRPK inhibitor), starting at 24 hours post-fertilisation (hpf). The media were changed daily. The embryos were raised until 120hpf and analysed for hyaloid structure.

Hyaloid experiment:

Tg(fli1:EGFP) zebra fish larvae were euthanized at 5 dpf in 0.003 % tricaine and fixed in 4% PFA/1xPBS overnight at 4°C. Fixed larvae were then washed in 1xPBS three times at RT for 20 min and incubated for 80 min at 37°C in 0.25% Trypsin/EDTA buffered with TRIS HCl (1.5 M,

pH 7.8) solution. Afterwards the larvae were washed three times and stored in PBS at 4°C until preparation. The larval hyaloid was then dissected under a stereoscope and displayed in PBS, in accordance with Jung’s protocol for visualization. Confocal images for phenotype evaluation were acquired using a confocal microscope (DM6000 B) with a scanner (TCS SP5 DS; Leica) using a 20x0.7 objective. The images were captured at 1024x 1024 pixels with 1,5µm Z-steps and 4.0 zoom factor.

### Site directed mutagenesis

Site directed S199 phospho-mutants of Phf6 were generated using the Q5® Site-Directed Mutagenesis Kit (New England Biolabs). Mutagenesis primers were generated using the NEBasechanger online calculator. The following primers were used (5′-3′):S199A: Forward-CCGAGATAGGGCGCCACACAGGA, Reverse-TAGGAGCTACTTTCCATTTCATCTG; S199E: Forward-CCGAGATAGGGAACCACACAGGAG, Reverse-TAGGAGCTACTTTCCATTTC. The PCR reactions were conducted with 1 X Q5 Hot Start High-Fidelity 2X Master Mix, 0.5 µM forwards and reverse primers and 10 ng/µl of template DNA (wild type Phf6 plasmid) were amplified for 25 cycles as per the manufacturer’s instructions. 1 µl of the PCR product was mixed with 5 µl of KLD reaction buffer, 1 µl of KLD enzyme mix and 3 µl of nuclease free water, and incubated for 5 minutes at room temperature. 5 µl of this mix was then transformed into DH5α competent cells, and plated on selective agar. Single colonies were subsequently expanded in liquid culture, and plasmid isolated using the NucleoBond Xtra Midi Kit (Macherey Nigel). Successful mutagenesis was validated by commercial sequencing (Eurofins).

### Immunofluorescence imaging

All immunofluorescence images were examined and captured with a DMi8 Leica microscope, unless stated otherwise (e.g. hyaloid imaging).

### RNA immunoprecipitation (RIP)

RIP was performed as in 11 with minor changes. Cells (four 10 cm plates for each replicate/condition) were washed with PBS and lysed in RIP lysis buffer (100 mM KCl, 5 mM MgCl_2_, 10 mM HEPES pH 7, 0.5% IGEPAL CA-630, 0.1 mM DTT, cOmplete™ EDTA-free Protease Inhibitor Cocktail, RNasIN Promega, Cat # N2511). Lysate was homogenised (through a 21-gauge needle), and the corresponding beads were prepared for IP as described above (Immunoprecipitation). Ten percent of lysate was removed as input, beads were washed twice in NT2 buffer (50 mM Tris-HCl pH 7.4, 150 mM NaCl, 5 mM MgCl2, 0.05% IGEPAL CA-630). Lysate was added to beads and incubated for 2 hours at 4 °C. The supernatant was removed and beads were washed five times in ice-cold NT2 buffer. RNA was eluted by proteinase K digestion (resuspension in 150 μL Proteinase K buffer (50 mM Tris-HCl pH 7.4, 150 mM NaCl, 5 mM MgCl2, 0.05% IGEPAL CA-630, 1% SDS, followed by 20 µl of proteinase K and incubation at 55 °C for 30 minutes alongside the digestion of matched inputs). 230 μl of NT2 buffer was added to supernatants, followed by phenol-chloroform separation in heavy lock tubes. The aqueous phase was purified and DNase digested using a Zymo RNA clean and concentrator kit according to the manufacturer’s instructions. Purified RNA was either sequenced or processed for qPCR, as detailed below.

### RNA sequencing analysis

RNA samples were denatured and mRNA was enriched using oligo (dT) magnetic beads. Enriched mRNA was subsequently fragmented, and cDNA was synthesised from RNA fragments using random primers, followed by second strand synthesis. Double-stranded cDNA fragments were subjected to end-repair, and then a single ‘A’ nucleotide was added to the 3’ ends of the blunt fragments. This was followed by adaptor ligation and amplification of the cDNA fragments.

The resulting libraries were assessed for quality, single-stranded PCR products were produced via denaturation and circularised. Single-stranded circular DNA molecules were replicated via rolling cycle amplification, and a DNA nanoballs (DNB) containing multiple copies of DNA generated. Sufficient quality DNBs were then loaded into patterned nanoarrays using a high-intensity DNA nanochip technique and sequenced through combinatorial Probe-Anchor Synthesis (cPAS) and sequenced using the DNBSEQ™ platform.

RNA was mapped and analysed using the RaNA-Seq platform (41). FASTQ files were first pre-processed using Fastp (42), and expression quantification was performed with Salmon (43).

Following quality assessment, differential expression analysis was performed using the DESeq2 tool (44).

For RIP sequencing data, analysis of enriched motif occurrence in FASTA files was performed using the Integrated System for Motif Activity Response Analysis (ISMARA) platform (45).

### Co-immunoprecipitation (Co-IP)

Cells (one 10 cm culture plate) were lysed in lysis buffer (50 mM Tris-HCl (pH 8), 120 mM NaCl, 1 mM EDTA, 1% Triton X-100 and 0.5 % IGEPAL CA-630). Phf6 complexes (each based on Myc-tagged construct overexpression) were captured using ChromoTek Myc-Trap® Magnetic Agarose and incubated at 4℃ for four hours. Lysate was removed from beads and beads were washed five times in Co-IP wash buffer (50 mM Tris-HCl (pH 8), 120 mM NaCl, 1 mM EDTA, 1% Triton X-100) prior to complex elution in 1 x Laemmli buffer prior to Western blot).

### qPCR

RNA (either from RIP or isolated using the NucleoSpin RNA extraction kit (Macherey-Nagel, for total RNA) was reverse transcribed using the Maxima H Minus First Strand cDNA Synthesis Kit (Thermo Fisher Scientific) to manufacturer’s instructions. qPCR was performed using the Maxima Sybr qPCR kit (Thermo Fisher Scientific). The relative target gene expression was normalised to *Gapdh* levels unless otherwise stated.

### Alpha-Fold Modelling

Predicted Phf6 structures and RNA interactions were modelled using the AlphaFold Server (AlphaFold3) (46). Phf6 structures were generated using the annotated NP_081918.1 FASTA sequence, with amino acid substitutions modelled and functional domains annotated as described in the corresponding Figures.

### Actinomycin D stability assay

Cells (MCECs) were cultured under the respective experimental conditions, and Actinomycin D (10 µg/mL) added for the indicated time periods prior to lysis, RNA extraction and qPCR as described.

### 3’UTR luciferase assay

The pGL3 luciferase vector (Promega) was digested with FseI and XbaI (corresponding to the 3’ end of the luciferase). The 3’UTR of Notch2 was amplified for DNA isolated from MCECs using the following primers (5′-3′): Forward: ggaaagatcgccgtgtaattAAGAGTCTGCCTCAGCCTTAG; Reverse: tgtctgctcgaagcggccggTGAGCTTTGACATGGAACC

The amplified product was gel purified and inserted into the pGL3 vector using the NEBuilder HiFi DNA Assembly Cloning Kit as per the manufacturer’s instructions. Successful insertion was validated by commercial sequencing (Eurofins).

Luciferase constructed was transfected into cells alongside the respective constructs indicated in the respective Figure. The relative luciferase activity was then assessed using the Nano-Glo™ Luciferase Assay according to the manufacturer’s instructions. Briefly, an equal volume of assay reagent was added to the media of cells, and luminescence measured following a five-minute incubation at room temperature.

### Schematic Images

Schematic images were created in BioRender.

### Statistical analysis

Data are shown throughout as mean ± SEM. Corresponding *n* numbers are reported in the respective Figure legends. Significance testing was performed with GraphPad Prism 7.

Statistical significance was assessed using an unpaired Students t-test (2 experimental groups) or a 1-way ANOVA with Tukey multiple comparison test (when comparison was made between 3 or more experimental groups).

## RESULTS

### VEGF-A stimulation drives widespread changes in the phosphorylation of the EC RNA interactome

To investigate the global effects of VEGF-A stimulation on RBP phosphorylation in ECs, mouse cardiac ECs (MCECs) were subjected to acute (15 minutes) or prolonged (24 hours) stimulation with VEGF-A. Protein-RNA interactions were UV-crosslinked and RNA interactome capture (oligo d(T) precipitation followed by high stringency washing) was used to isolate the proteins in direct interaction with mRNA at the time of cross-linking (i.e. the ‘global’ mRNA bound proteome). Isolated RBPs and phospho-sites were then identified by side-by-side quantitative total proteomic and phosphoproteomic analysis (Fig. 1A)

**Figure 1.**
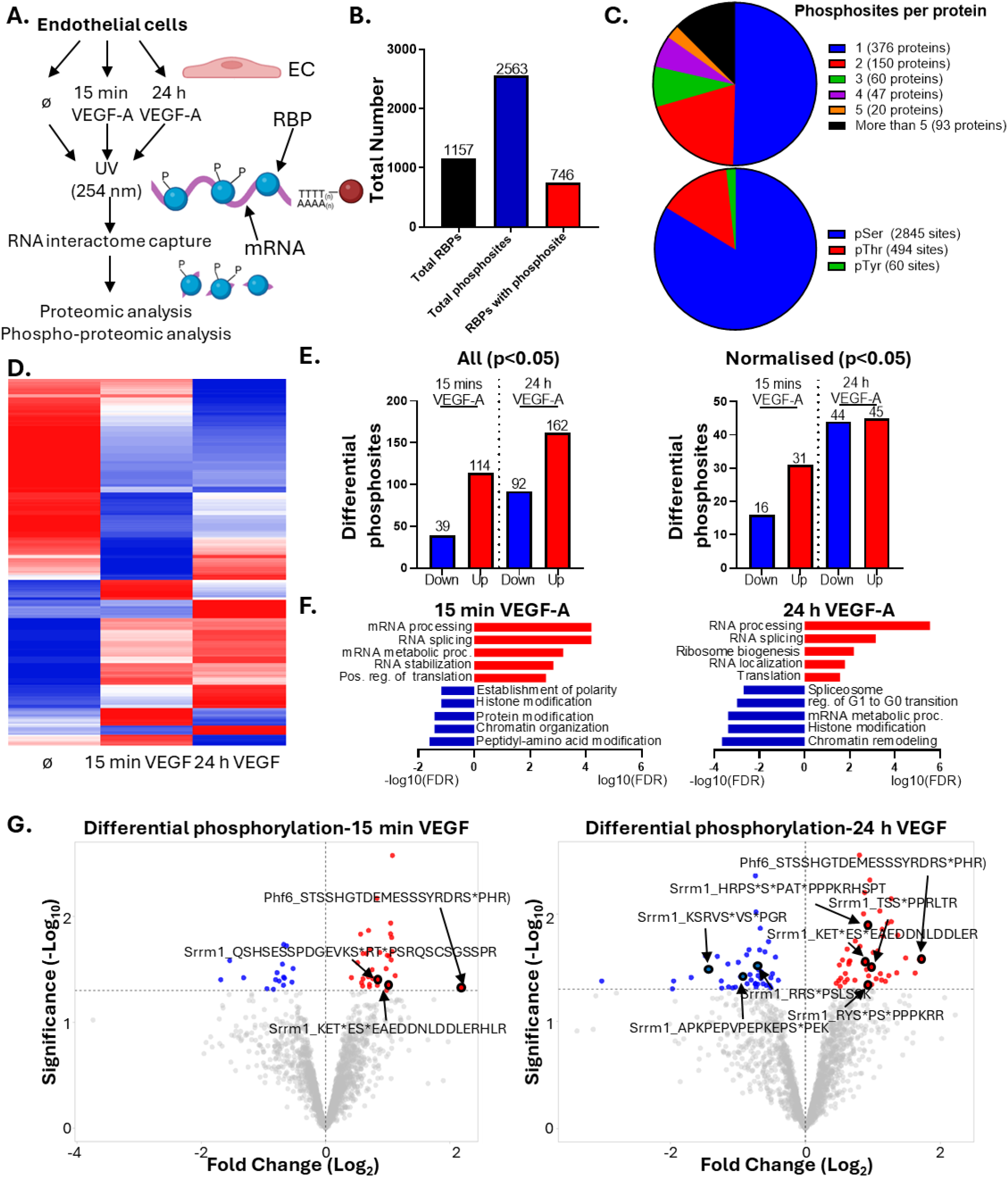
Identification of global VEGF-A driven changes in phosphorylation of the mRNA bound proteome in MCECs. A Schematic overview of experimental approach. B. Total number of RBPs identified in proteomic analysis of RIC isolates and phosphorylation sites identified in phosphoproteomic analysis of RIC isolates. C. Number of phosphorylation sites contained within each identified RBP, and proportion of phosphorylation occurring at Ser, Thr and/or Tyr. D. Heatmap of differential changes in RBP phosphorylation occurring after 15 minutes or 24 hours of VEGF-A stimulation (averaged intensities from three replicates per condition, columns ordered by Spearman rank correlation, and rows were coloured by Z score). E. Number of differential phosphosites (P<0.05) within RBPs after 15 minutes or 24 hours of VEGF-A stimulation, shown in both the presence and absence of the RBP abundance in the corresponding RIC proteome (i.e. changes in the abundance of the protein bound to RNA). F. Enriched GO biological process terms of differentially (P<0.05, normalised to RBP abundance in RIC proteome) phosphorylated RBPs after 15 minutes or 24 hours of VEGF-A stimulation (upregulated shown at Log_10_FDR, downregulated as - Log_10_FDR. G. Volcano plots of differential phosphosites (P<0.05) after 15 minutes or 24 hours of VEGF-A stimulation, differential phosphorylation sites within Phf6 and Srrm1, chosen for subsequent validation, are labelled. RIC; RNA interactome capture.

This identified a total of 1157 RBPs and 2563 phosphosites, with 746 (approximately two-thirds) of the identified RBPs containing at least one phosphosite irrespective of VEGF-A stimulation (Fig. 1B, Sup. Fig. 1A). As validation of the specificity of our interactome capture approach, analysis of enriched GO biological process and molecular function terms in the identified RBPs showed a clear enrichment for RNA-binding related functions (Sup. Fig. 1B). The majority of RBPs contained only one or two phosphosites, but we also identified a number of ‘hyper’-phosphorylated RBPs containing up to 163 phosphosites, with the majority of phosphorylation occurring at serine resides (Fig. 1C, Sup Fig. 1C, D).

Analysis of the changes in RBP phosphorylation after 15 minutes or 24 hours of VEGF-A stimulation found highly dynamic changes, with distinct phosphorylation and dephosphorylation of RBPs occurring in both acute and prolonged stimulation (Fig. 1D). When focussing on the significantly changed differential phosphosites, we found 114 upregulated and 39 downregulated after 15 minutes and 162 upregulated and 92 downregulated at 24 hours of stimulation, which decreased after normalisation to changes in abundance of the respective RNA bound RBPs showing that distinct subsets of RBPs change in their global or specific interactions with RNA upon phosphorylation (Fig. 1E., Sup. Fig. 1E, F).

Analysis of enriched GO biological process terms among differentially phosphorylated RBPs, after normalisation, indicated that differential phosphorylation occurs in RBPs involved in the widespread coordination of RNA metabolism including splicing, RNA stability, translation and RNA processing, suggesting that RBP phosphorylation downstream of VEGF-A results in an orchestrated response by the post-transcriptional machinery (Fig. 1F). This was predicted to be most likely downstream of canonical effector kinases such as AKT, MAPK and MTOR (Sup. Fig. 1G). We visualised the most pronounced differential phosphosites, finding multiple differential phosphosites within Serine and Arginine Repetitive Matrix 1 (Srrm1) that were significantly altered, whereas one particular site within PHD Finger Protein 6 (Phf6) at serine 199 (S199) showed the strongest increase in phosphorylation after both 15 minutes and 24 hours of VEGF-A stimulation (Fig. 1G).

### VEGF-A increases phosphorylation and RNA binding of Srrm1 and Phf6

We therefore chose Phf6 and Srrm1 for subsequent analysis, first validating the interaction of both proteins with mRNA (Sup. Fig. 2A). Overexpression of tagged Srrm1 or Phf6 constructs, followed by 15 minutes or 24 hours of VEGF-A stimulation in mouse ECs, immuno-precipitation and Western blot analysis confirmed increased serine phosphorylation in Srrm1 after 24 hours of VEGF-A stimulation, and increased serine phosphorylation within Phf6 after both 15 minutes and 24 hours of phosphorylation in protein purified from total lysate as well as protein purified from isolated RNA binding proteins (IP following RIC) (Sup. Fig. 2A).

**Figure 2.**
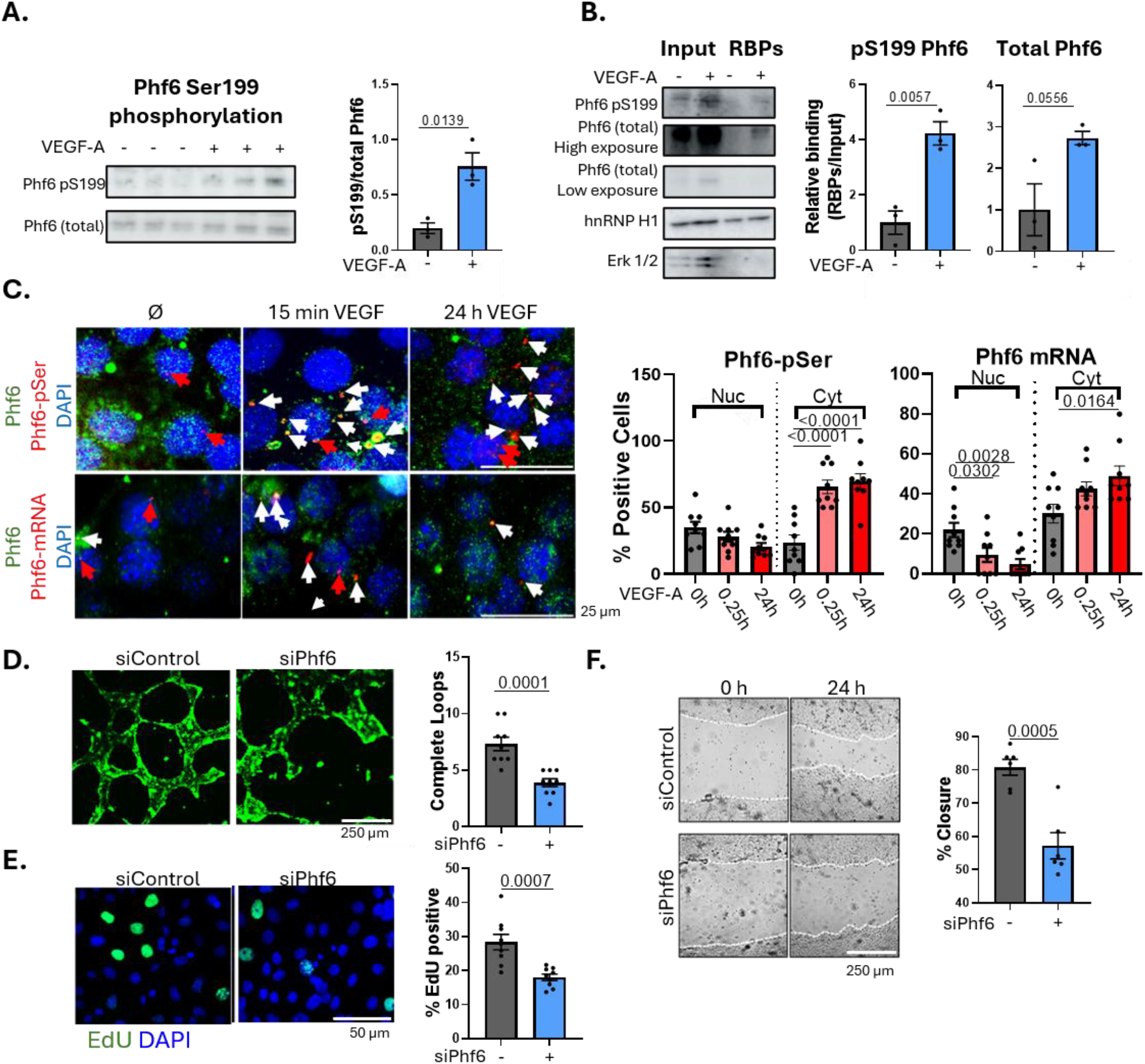
Phf6 is differentially phosphorylated after VEGF-A stimulation and contributes to changes in EC phenotype. A. Validation of Phf6 S199 phosphorylation upon VEGF-A stimulation. MCECs were stimulated with VEGF-A (10 ng/ml) for 24 hours followed by lysis and Western blot using a phospho-S199 Phf6 specific antibody (followed by total Phf6). Quantifications reflect the relative change in S199 phosphorylation (relative to total Phf6). Data shown as mean ± SEM, significance assessed by t-test, significance indicated. B. Validation of VEGF-A driven changes in S199 phosphorylation in RNA bound Phf6. MCECs were stimulated with VEGF-A for 24 hours followed by UV-crosslinking, RNA interactome capture and Western blot using a phospho-S199 Phf6 specific antibody (followed by total Phf6). Quantifications reflect the relative change in the abundance of either pS199 Phf6 or total Phf6 in RNA interactome capture eluates upon VEGF-A stimulation relative to input. Data shown as mean ± SEM, significance assessed by t-test, significance indicated. C. Visualisation of VEGF-A driven changes in the phosphorylation and mRNA interactions of Phf6. A proximity ligation assay (PLA) in MCECs stimulated with either 15 minutes or 24 hours of VEGF-A stimulation (10 ng/ml) using an antibody recognising endogenous Phf6, along with either a p-Ser probe or biotinylated oligo d(T) probe followed by anti-biotin antibody, followed my immunofluorescence microscopy, was to visualise changes in the phosphorylation and RNA interactions of Phf6 in MCECs following 15 minutes or 24 hours of VEGF-A stimulation. PLA signal (phosphorylated Phf6 or Phf6 –mRNA interactions) is shown in red, total Phf6 shown in green (DAPI in blue). White arrows show examples of cytoplasmic PLA signal and red arrows show examples of nuclear PLA signal. Quantifications show percentage of cells per image positive for nuclear and/or cytoplasmic PLA signal (n=9, significance assessed by one-way ANOVA, significance indicated). D. Effects of Phf6 knockdown (siRNA, 48 hours, ± 10 ng/ml VEGF-A stimulation for 24 h) on tube formation in MCECs plated in Matrigel for 24 hours. Tubes were fluorescently labelled with BCECF AM Ester prior to imaging. Representative images and quantifications of the number of complete loops per visible field, n=3 triplicates/condition, data shown as average ±SEM significance assessed by t-test, significance indicated. E. Effects of Phf6 knockdown (siRNA, 48 hours, ± 10 ng/ml VEGF-A stimulation for 24 h) on EdU incorporation (proliferation) in MCECs. Representative images and quantifications of EdU incorporation, n=3 triplicates/condition, quantified as percentage EdU positive cells per image data shown as average ±SEM significance assessed by t-test, significance indicated. F. Effects of Phf6 knockdown (siRNA, 48 hours, ± 10 ng/ml VEGF-A stimulation for 24 h) on MCEC migration as assessed by scratch wound assay. Representative images and quantification of the percentage closure 24 hours post-scratch, n=3 triplicates/condition. Data shown as average ±SEM significance assessed by t-test, significance indicated.

Using an antibody specific to Phf6-pSer199, we validated a specific increase in Ser199 phosphorylation following VEGF-A stimulation in total protein lysate. Further to this, analysis of the relative change in abundance of Phf6 and Phf6-pSer199 in RNA interactome capture eluates (i.e. RNA bound) following VEGF-A stimulation revealed a close correlation between S199 phosphorylation and a VEGF-A driven increase in the RNA binding of Phf6 (Fig. 2B).

Interestingly, this also showed that there is a particular enrichment of Phf6 with Ser199 phosphorylation compared to total Phf6 in RIC isolates, again suggesting a link between S199 phosphorylation and the RNA binding of Phf6.

Next, we performed a proximity ligation assay (PLA) in MCECs following 15 minutes or 24 hours of VEGF-A stimulation using an antibody specific to endogenous Phf6 together with either a phospho-serine specific probe to detect and visualise the endogenous Phf6 phosphorylation or a biotinylated oligo d(T) probe and anti-biotin antibody to detect and visualise Phf6-mRNA interactions. This further confirmed an increase both the phosphorylation and RNA binding of Phf6 upon VEGF-A stimulation, occurring primarily in the cytoplasm (Fig. 2C). We found a similar increase in the cytoplasmic phosphorylation and RNA binding of Srrm1 (Sup. Fig. 2C). This correlated with a relative shift of either protein from the nucleus to the cytoplasm upon VEGF-A stimulation (Sup. Fig. 2D).

### Differentially phosphorylated RBPs, Srrm1 and Phf6, regulate angiogenic phenotypes in murine ECs

Given the significant changes in the phosphorylation of RNA bound Phf6 and Srrm1 following VEGF-A stimulation, we hypothesised that changes in the RNA binding of these proteins play roles in coordinating EC responses to VEGF-A with regard to angiogenic behaviour. We knocked-down Phf6 or Srrm1 in MCECs and investigated the effects on various phenotypic characteristics associated with enhanced angiogenesis. Knock-down efficiency was confirmed by qPCR (Sup. Fig. 3A). We found that Phf6 knockdown increased the capacity of MCECs to form tubes on Matrigel (Fig. 2D). Similarly, Phf6 knockdown significantly inhibited proliferation and migration in MCECs when compared to control cells, together suggesting that Phf6 plays a proangiogenic role (Fig. 2 E and F). Srrm1 knockdown also inhibited tube formation and migration, although increasing proliferation in MCECs (Sup. Fig. 3 B-D).

**Figure 3.**
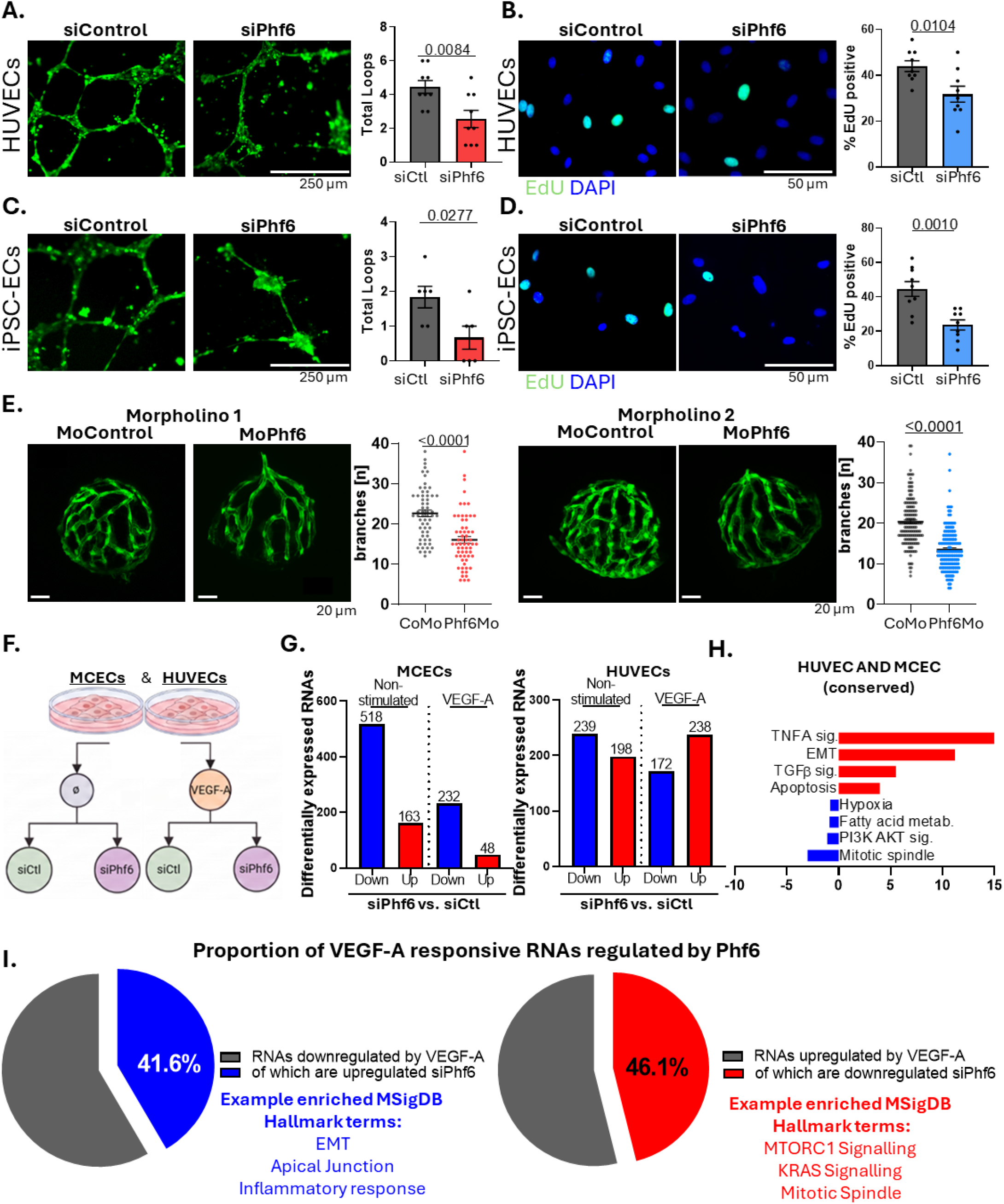
The role of Phf6 in VEGF-A driven angiogenesis. A and B. Effects of Phf6 knockdown on tube formation and proliferation in HUVECs. Phf6 was knocked down (siPhf6) in HUVECs for 48 hours, followed by plating of cells onto Matrigel and assessment of tube formation (A) or assessment of proliferation by an EdU incorporation assay (B). Representative images , n=3 triplicates/condition, data shown as average ±SEM significance assessed by one-way ANOVA, significance indicated. C and D. Effects of Phf6 knockdown on tube formation and proliferation in iPSC derived ECs. Phf6 was knocked down (siPhf6) in iPSC derived ECs for 48 hours, followed by plating of cells onto Matrigel and assessment of tube formation (C) or assessment of proliferation by an EdU incorporation assay (D). Representative images , n=3 duplicates/condition, data shown as average ±SEM significance assessed by one-way ANOVA, significance indicated. E. Hyaloid vasculature in zebrafish (fli1EGFP) 5 dpf following morpholino based Phf6 knockdown (both 6 ng/nl, as well as control morpholino) using two independent Phf6 targeting morpholinos. For the morpholino 1 experiment n= control morpholino: 61, morpholino 1: 62, for the morpholino 2 experiment n= control morpholino: 151, morpholino 2:147. Significance was assessed by t-test, error bars shown average ±SEM. F. Schematic overview of RNA-sequencing approach. Phf6 was knocked down (siPhf6, 48 h) in MCECs or HUVECs in the presence and absence of VEGF-A stimulation (24 h) followed by RNA extraction and RNA sequencing. G. Number of differentially expressed RNAs in MCECs or HUVECs following Phf6 knock-down (P < 0.05, Log2 FC < 0.5). H. Enriched Msig DB Hallmark terms of conserved (i.e. up- or downregulated in both MCECs and HUVECs) differentially expressed RNAs (up-regulated in both shown in red, downregulated in both down as blue). I. The proportion of overlap of RNAs that were differentially expressed (P<0.05) in HUVECs following VEGF-A stimulation (24 h) that showed opposing changes in expression following Phf6 knockdown (siPhf6, 48 h), with example enriched Msig DB Hallmark terms for each shown.

### Phf6 promotes angiogenic features in human ECs

Based on this consistent phenotype as well as the significant increase in phosphorylation observed at the single S199 site, we then focussed on Phf6 for subsequent further characterisation. To investigate if the effects of Phf6 are also conserved in human ECs, we knocked down Phf6 in both primary HUVECs (human umbilical vein endothelial cells) and human iPSC (induced pluripotent stem cell) derived ECs. We found that Phf6 knockdown significantly inhibited tube formation and reduced proliferation in both human EC types (Fig. 3A-D).

### Knock-down of Phf6 in zebrafish reduces hyaloid vasculature formation

Given the effects of Phf6 on angiogenic like properties in ECs *in vitro,* we investigated the effects of Phf6 knock-down on vascular development *in vivo* in zebrafish using a morpholino based approach. We used two independent morpholinos, morpholino 1 (MO) utilising a splice blocking approach and morpholino 2 utilising a translation blocking approach to knockdown Phf6 in zebrafish embryos. The efficacy of both approaches was validated by PCR or Western blot (Sup. Fig. 3E). Assessment of hyaloid vasculature formation at 5 dpf (days post-fertilisation) revealed that loss of Phf6 significantly inhibits vascular branching (Fig. 3E).

### Loss of Phf6 results in the differential expression of angiogenesis related RNAs in mouse and primary human ECs

For deeper insight into how Phf6 coordinates changes in the endothelial response to VEGF-A stimulation, we used RNA sequencing to identify differential changes in RNA expression in MCECs or HUVECs following Phf6 knockdown in the presence and absence of VEGF-A stimulation (Fig 3F).

We investigated differential RNA expression after Phf6 knockdown in MCECs. In both non-stimulated and VEGF-A stimulated MCECs, knockdown of Phf6 resulted primarily in the decreased expression of RNAs, with the effects more pronounced in non-stimulated cells (Fig. 3G, Sup. Fig. 4A). To understand the downstream consequences of Phf6-driven changes in RNA expression, we identified the enriched MSigDB (Molecular Signatures Database) hallmark terms for differentially expressed RNAs. This found, for instance, an upregulation of RNAs involved in epithelial-mesenchymal transition (EMT), TNFα and TGFβ signalling and apoptosis, as well as a downregulation of RNAs related to angiogenesis, KRAS and Hedgehog signalling and hypoxia response (Sup. Fig. 4B).

**Figure 4.**
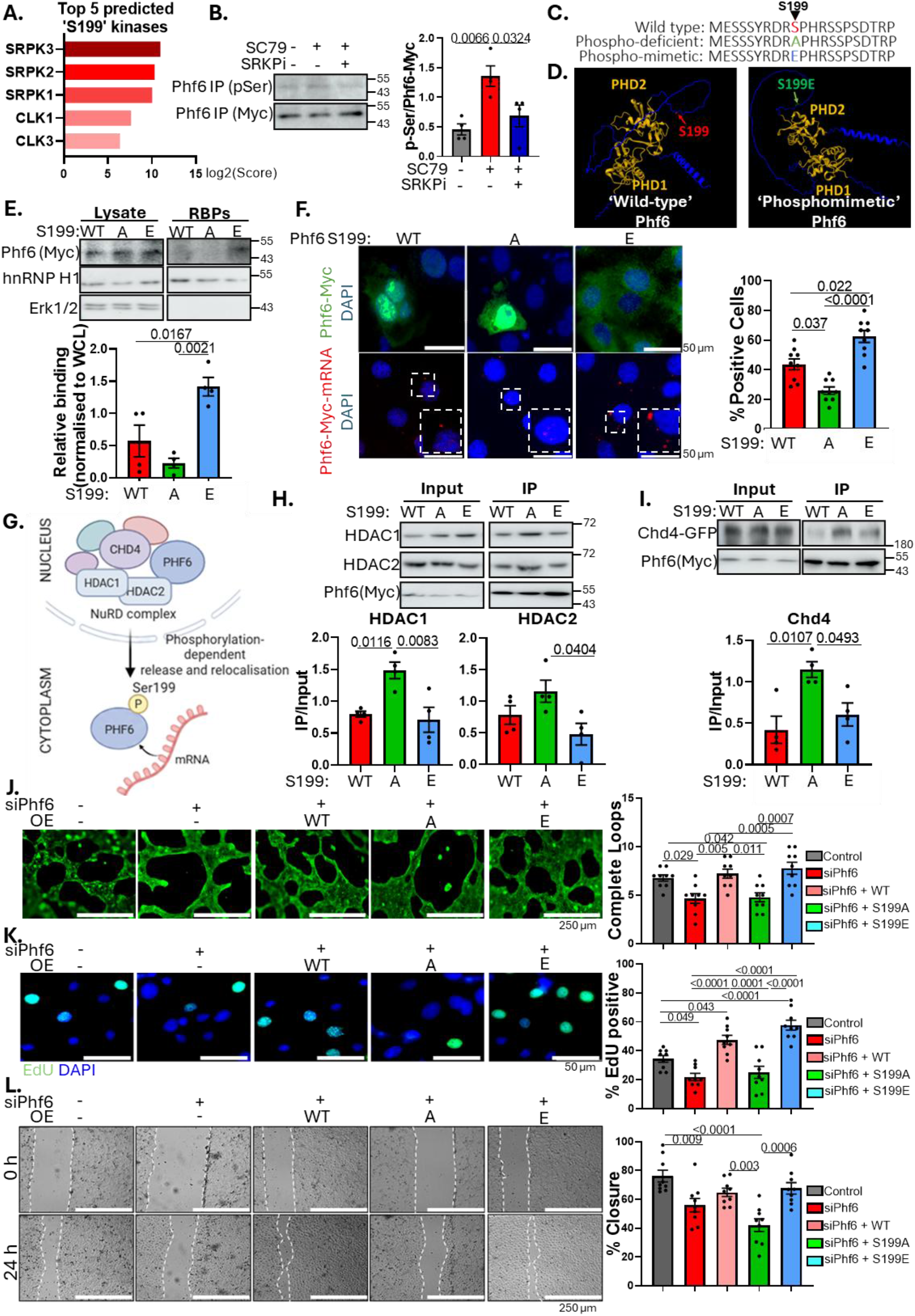
Phf6 phosphorylation at Ser199 regulates angiogenic phenotype. A. The top five predicted kinases upstream of Phf6 Ser199 phosphorylation, using the score site function of The Kinase Library. B. The effects of AKT and SRPK on Phf6 phosphorylation. Phf6 was overexpressed in MCECs for 48 hours. SC79 (10 µM, 24 hours) was used as a stimulator of AKT and MSC1186 (SRPKi, 10 nM 24 hours) as an inhibitor of SRPK, followed by Phf6 IP (anti-Myc) and Western blot analysis of total serine phosphorylation. Quantifications show average ±SEM, n=4, significance assessed by one-way ANOVA, significance indicated.C. Site directed mutagenesis was used to generate an Ser199 phosphodeficient mimic Phf6 by S199A substitution and a S199 phoshomimetic Phf6 by S199E substitution. D. Alpha-fold structure predicted of wild type and phosphomimetic Phf6 (S199E). The Phf6 PHD domains are highlighted in yellow. E. RNA interactome capture following 48 h construct overexpression and Western blot analysis was used to compare the relative expression and RNA binding of wild-type (WT), phosphodeficient (S199A) and phosphomimetic (S199E) Phf6 constructs. Quantifications show average ±SEM, n=4, significance assessed by one-way ANOVA, significance indicated. F. Immunofluorescence microscopy of MCECs following 48 hours of wild-type (WT), phosphodeficient (S199A) or phosphomimetic (S199E) Phf6 overexpression. Top-anti-myc staining of Phf6 localisation (green), bottom-PLA of Phf6-mRNA interactions using a biotinylated oligo d(T) probe and anti-myc and anti-biotin antibodies. Phf6-mRNA interactions shown in red. G. Schematic overview of proposed effects of Ser199 phosphorylation on the interaction between Phf6 and nuclear complex (i.e. NuRD complex binding).H. The effects of S199 phosphorylation on the interaction between Phf6 and HDAC1 and HDAC2. Wild-type (WT), phosphodeficient (S199A) or phosphomimetic (S199E) (all Myc tagged) Phf6 were overexpressed in MCECs for 48 hours, followed by co-IP (Myc-trap) and Western blot analysis of the interaction with HDAC1 and HDAC2. N=3, quantifications show average ±SEM, significance assessed by one-way ANOVA, significance indicated. I. The effects of S199 phosphorylation on the interaction between Phf6 and Chd4. Wild-type (WT), phosphodeficient (S199A) or phosphomimetic (S199E) (all Myc tagged)Phf6 were overexpressed along with Chd4 (GFP-tagged) in MCECs for 48 hours, followed by co-IP (Myc-trap) and Western blot analysis of the interaction with Chd4. N=3, quantifications show average ±SEM, significance assessed by one-way ANOVA, significance indicated. J,K and L. Phf6 was knocked down in MCECs (siPhf6, 48 h) alongside co-transfection of either wild-type (WT), phosphodeficient (S199A) or phosphomimetic (S199E) Phf6 constructs, followed by plating of cells onto Matrigel and assessment of tube formation (I), assessment of proliferation by an EdU incorporation assay (J) or assessment of migration by a 24 h scratch assay (K). Representative images. Each was done three times in triplicate, quantifications show average ±SEM, significance assessed by one-way ANOVA, significance indicated.

Phf6 knockdown in HUVECs similarly resulted in altered RNA expression, also including an upregulation of RNAs involved in TNFα signalling, EMT and apoptosis and a downregulation of RNAs involved in mitosis, cholesterol homeostasis and apical junctions, for example (Sup. Fig. 4C and D). Overlap of conserved differentially expressed RNAs (RNAs that were either up-or downregulated in both MCECs and HUVECs) revealed an upregulation of RNAs involved in pro-inflammatory (e.g. TNFα) and pro-mesenchymal processes (e.g. epithelial-mesenchymal transition and TGF-β signalling) and a down regulation of RNAs involved in mitosis, metabolism, hypoxia and AKT signalling (Fig. 3H, Sup. Fig. 4E).

### Phf6 regulates VEGF-A driven transcriptomic changes

We next wanted to elucidate the overall extent to which Phf6 is responsible for mediating VEGF-A driven changes in RNA expression. VEGF-A stimulation alone (siCtl VEGF-A vs. siCtl non-stimulated) led to increased expression of RNAs associated with mitosis and E2F factors, MTORC1 and cholesterol homoeostasis, as well as a downregulation of RNAs related to glycolysis, hypoxia, apical junctions and EMT (Sup. Fig. 5A and B). To identify which part of the VEGF-A mediated transcriptomic response is regulated by Phf6, we focussed on RNAs down-regulated by VEGF-A stimulation and up-regulated by Phf6 knockdown, and *vice versa*. Interestingly, we found a very large proportion (41.6%) of RNAs that were significantly downregulated by VEGF-A and were upregulated by siPhf6. 46.1% of RNAs upregulated by VEGF-A were downregulated by siPhf6 during VEGF-A stimulation, demonstrating the importance of Phf6 in mediating VEGF-A driven changes in RNA expression (Fig. 3I, Sup. Fig. 5C and D). Based on our results, we hypothesise that Phf6 facilitates a decrease in the expression of RNAs involved in EMT and apical junctions and promotes the expression of RNAs involved in MTORC1 signalling, KRAS signalling and mitosis to promote EC proliferation, sprouting and angiogenesis (Sup. Fig. 5E).

**Figure 5.**
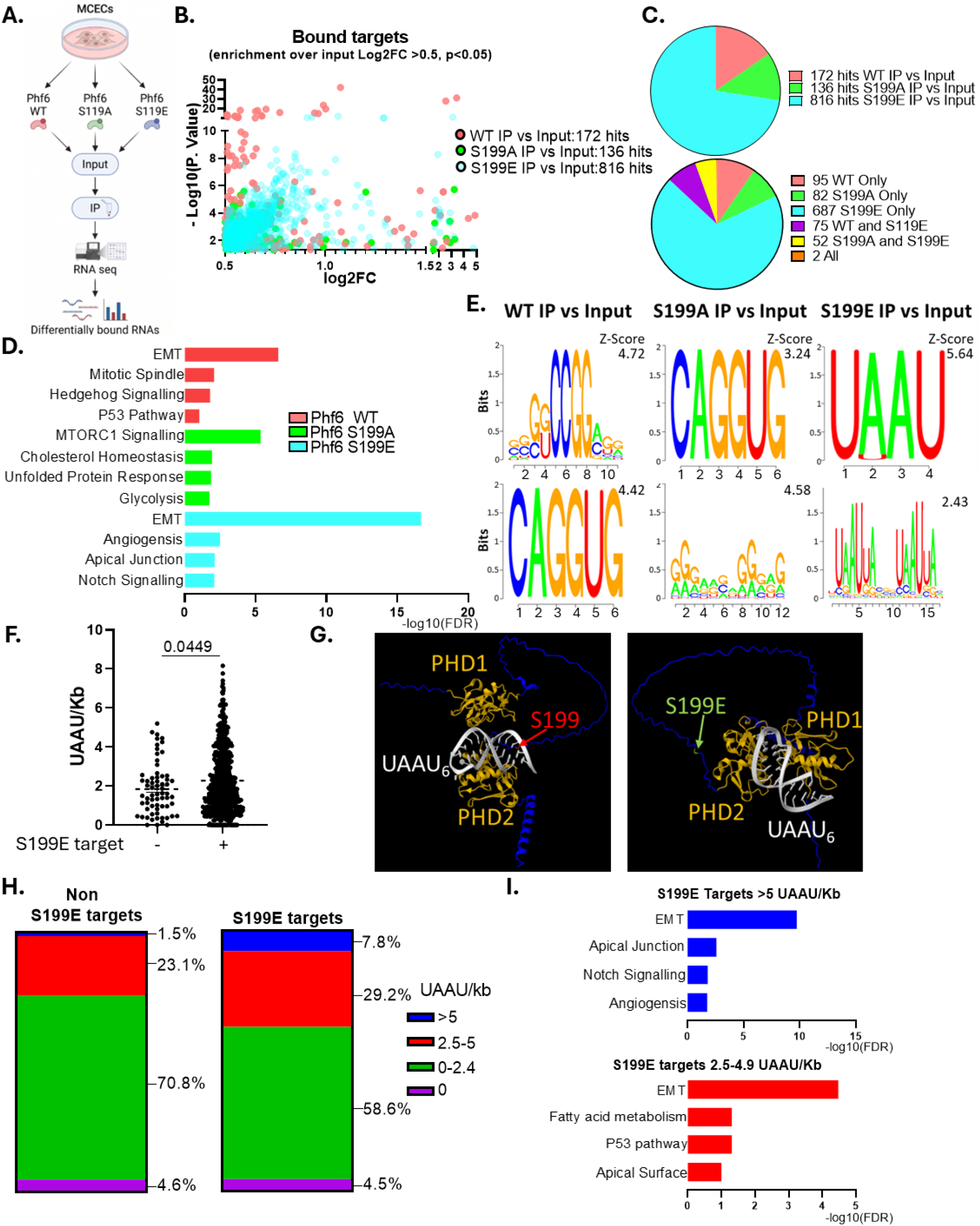
Phf6 phosphorylation at Ser199 drives differential RNA binding. A. Schematic overview of experimental design to identify the differential changes in RNA interactions of wild-type (WT), phosphodeficient (S199A) and phosphomimetic (S199E) Phf6 using RNA immunoprecipitation and RNA sequencing (RIP-seq.). B. Significantly enriched target RNAs (IP vs Input, Log2FC >2, p<0.05) identified in wild type (WT), phospho-deficient (S199A) or phosphomimetic (S199E) Phf6 IPs. C. Top: Visualisation of the total number of target RNAs (hits) of WT, S199A or S199E Phf6 (identified in fig. 4B), and Bottom: The distribution of these targets across the conditions. D. Enriched Msig DB Hallmark terms of the target RNAs bound to wild-type (WT), phosphodeficient (S199A) and phosphomimetic (S199E) Phf6. E. The individual enriched peaks in WT, S199A or S199E Phf6 IP vs inputs were scanned for differential motif enrichments. Significantly enriched motifs across the three conditions are shown, with the individual z-score annotated. F. The target RNAs identified as bound by phosphorylated Phf6 (S199E bound) were compared for the frequency of UAAU motifs against targets not-bound by phosphorylated Phf6 (i.e. all other targets), normalised by transcript length. Significance was assessed by an unpaired t-test. G. Alpha-fold predicted modelling of the interaction between with wild type Phf6 (S199 highlighted in red, left), or phosphomimetic Phf6 (S199E highlighted in green, right) with UAAU RNA motifs. H. The proportion of S199E targets and non-S199E targets containing varying frequencies of UAAU motifs. I. Msig DB Hallmark terms of ‘hyper’ UAAU rich (> 5 UAAU/Kb, shown in blue) and ‘high’ UAAU rich (2.5-4.9 UAAU/Kb, shown in red) S199E targets.

### Phosphorylation of Phf6 at Ser199 occurs downstream of SRPK

We then wanted to analyse the importance of Ser199 (S199) phosphorylation within Phf6 in mediating its proangiogenic function. We first tested the effects of inhibiting Erk (Extracellular signal-regulated kinase)1/2 or AKT (Protein kinase B), both known to mediate VEGF-A signals within ECs, on Phf6 phosphorylation by overexpressing Phf6 and stimulating MCECs with VEGF-A in the presence of pharmacological inhibitors (PD 0325901 as Erk inhibitor and Pictilisib as AKT inhibitor) of either kinase, followed by IP and Western blot for total serine phosphorylation (Sup. Fig. 6A). Only AKT inhibition, but not Erk1/2 inhibition resulted in a significant decrease in Phf6 phosphorylation, suggesting AKT as a likely upstream kinase. Predictive analysis using the Kinase library Score Site function (47) suggested Serine-arginine protein kinases 1, 2 and 3 (SRPK1, SRPK2 and SRPK3) as potential upstream kinases of S199 phosphorylation (Fig. 4A). Indeed, we found a dose-dependent decrease in Phf6 phosphorylation in response to a pan-SRPPK inhibitor (Sup. Fig. 6B). We hypothesised that VEGF-A stimulation activates AKT, which subsequently activates SRPK that in turn phosphorylates Phf6 at S199. We confirmed that AKT activation (using SC79, a selective AKT agonist) led to significantly increased Phf6 phosphorylation (in the absence of VEGF-A), which was inhibited by SRPK inhibition (Fig. 4B). To further support Phf6 as downstream mediator of AKT, we found that knock-down of Phf6 in HUVECs with constitutive AKT overexpression inhibited the AKT driven increase in proliferation and tube formation (Sup. Fig. 7 A and B). Interestingly, we found that inhibition of SRPK in zebrafish, by addition of MSC-1186 to the water at 24 hpf significantly inhibited vascular branching of the hyaloid vessels, whilst not effecting mortality, in a similar manner as Phf6 morpholinos (Sup. Fig. 8). Together this suggests that SRPK kinases, downstream of AKT, may be responsible for Phf6 phosphorylation at Ser199 and are, in turn, upstream mediators of Phf6 driven changes in the angiogenic phenotype.

**Figure 6.**
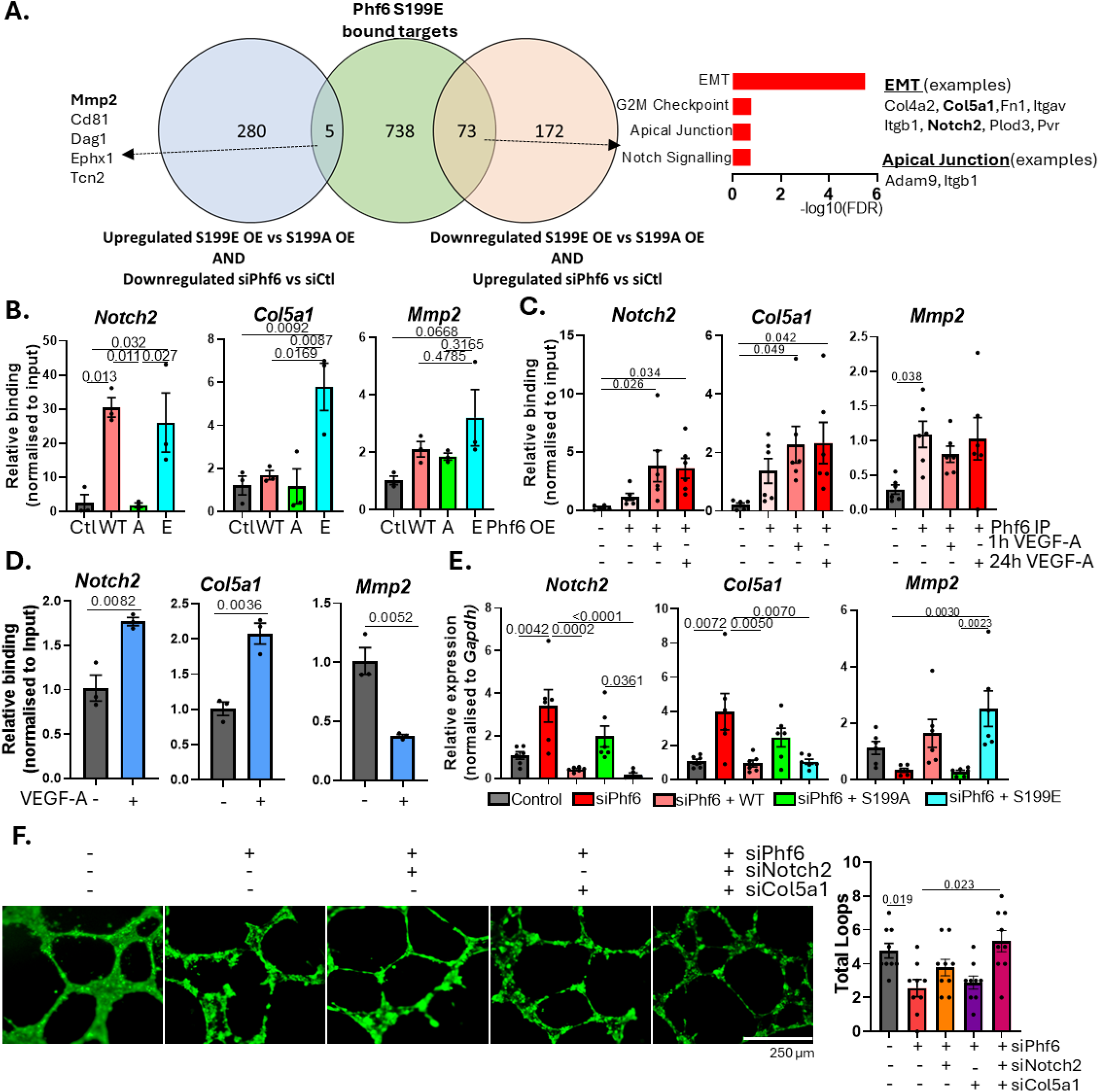
Phf6 differentially binds to regulates angiogenesis related RNAs. A. Overlap of RNAs that are targets of phosphorylated Phf6 and are either up-regulated (downregulated by siPhf6 and upregulated by S199E overexpression compared to S199A overexpression) or downregulated (upregulated by siPhf6 and downregulated by S199E overexpression compared to S199A overexpression). Enriched M Sig DB Hallmark terms for RNAs bound by and downregulated by phopshorlyated Phf6. B. RIP qPCR validation of selected targets of Phf6. Wild type (WT), phospho-deficient (S199A) or phosphomimetic (S199E) Phf6 was overexpressed in MCECs followed by immunoprecipitation of Phf6 constructs (anti-Myc) and qPCR analysis of the enrichment of selected target RNAs, normalised to expression in input lysates. Ctl indicates empty Myc vector. N=3. Data shown as average ±SEM significance assessed by one-way ANOVA, significance indicated. C. MCECs were stimulated with VEGF-A (10 ng/ml) for 0 hours, 1 hour or 24 hours, followed by immunoprecipitation of endogenous Phf6 and qPCR analysis of the enrichment of selected target RNAs compared to expression in input lysate. Phf6 IP – represents IgG control. . N=6. Data shown as average ±SEM significance assessed by one-way ANOVA, significance indicated. D. HUVECs were stimulated in the presence and absence of VEGF-A (10 ng/ml for 24 h), followed by immunoprecipitation of endogenous Phf6 and qPCR analysis of the enrichment of selected target RNAs compared to expression in input lysate. N=3. Data shown as average ±SEM significance assessed by students t-test, significance indicated. E. Phf6 was knocked down (siPhf6) in MCECs for 48 hours followed by 24 hour rescue with wild type (WT), phospho-deficient (S199A/A) or phosphomimetic (S199E/E) Phf6 for 24 hours, followed RNA isolated and qPCR analysis of the relative expression of selected target RNAs of Phf6 (normalised to *Gapdh).* N=6. Data shown as average ±SEM significance assessed by one-way ANOVA, significance indicated. F. Effects of Phf6 and simultaneous Notch2 or Col5a1 knockdown (siRNA, 48 hours, +10 ng/ml VEGF-A stimulation for 24 h) on tube formation in MCECs plated in Matrigel for 24 hours. Representative images and quantifications of the number of complete loops per visible field, n=3 triplicates/condition, data shown as average ±SEM significance assessed by one-way ANOVA, significance indicated.

**Figure 7.**
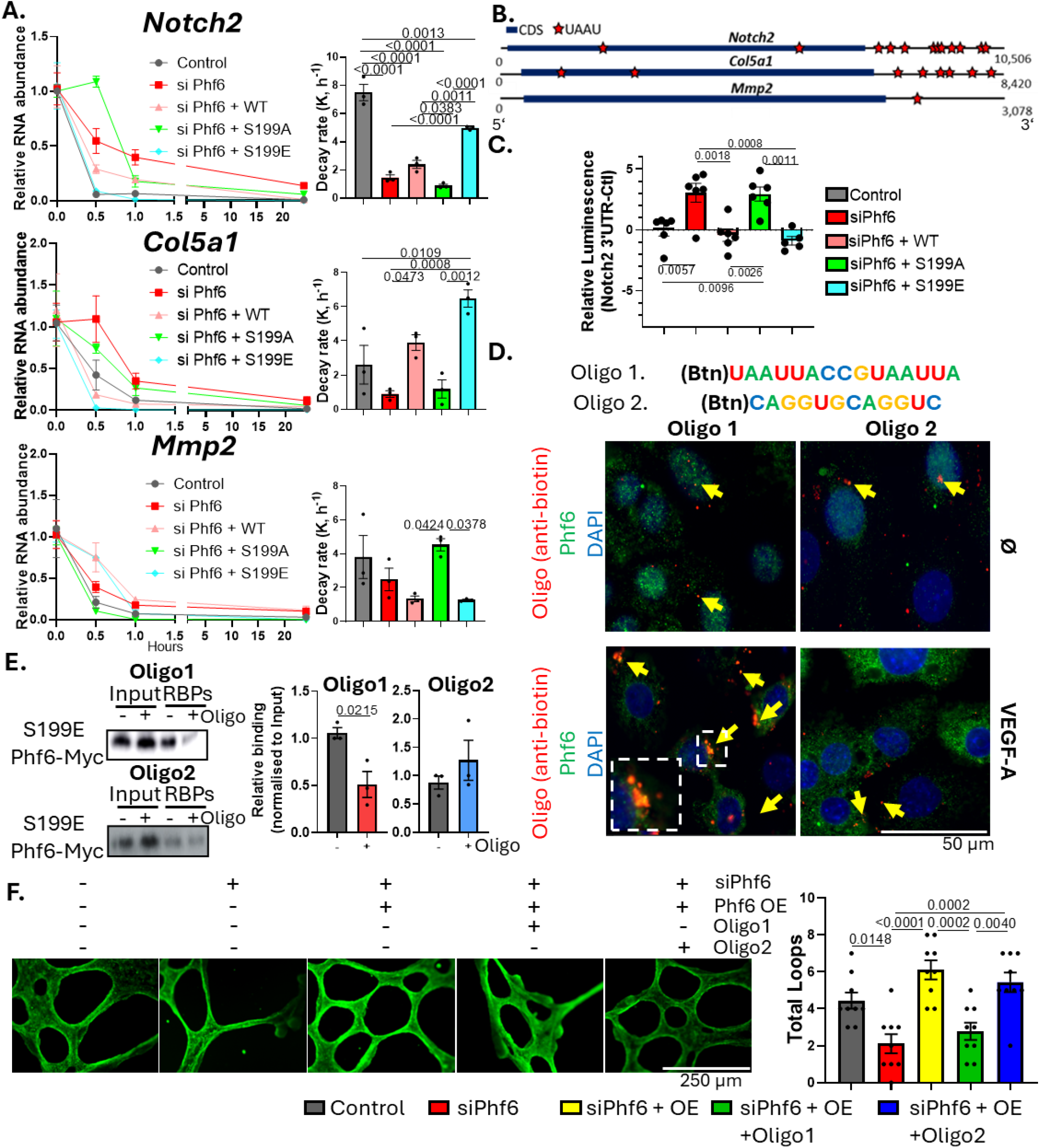
Phopshorylation driven in Phf6-RNA interactions differentially coordinates target RNA stability. A. The effects of Phf6 phosphorylation on target RNA stability. Phf6 was knocked down in MCECs (siPhf6 48 hours) alongside overexpression of wild type (WT), phospho-deficient (S199A) or phosphomimetic (S199E) Phf6. A time-course of actinomycin D was added to inhibit de novo transcription. qPCR was used to assess the relative abundance of selected Phf6 targets at each timepoint, and the decay rate was inferred from the respective curves. N=3, data shown as average ±SEM, significance assessed by one-way ANOVA, significance indicated. B. Schematic overview of the relative abundance of UAAU motifs within selected Phf6 targets. C. The 3’UTR of Notch2 was cloned downstream of a luciferase reporter system. The reporter construct was transfected into cells alongside siPhf6 together with either WT Phf6, S199A or S199E overexpression. Reporter stability was subsequently assessed by measurement of luciferase activity. N=6, data shown as average ±SEM, significance assessed by one-way ANOVA, significance indicated. D. Competitive inhibitor oligonucleotides were designed based on the differential binding preference of phosphorylated/unphosphorylated Phf6 (Oligo1 based on motifs preferentially bound by S199E Phf6, and Oligo2 based on motifs preferentially bound by WT and S199A Phf6). Oligos were designed as 2’O-Methyl RNA modified oligos incorporating a biotin tag at the 5’ end. Immunofluorescence staining following oligonucleotide transfection was used to visualise interactions between endogenous Phf6 and oligos in the presence and absence of VEGF-A stimulation (10 ng/ml for 24 hours), with examples of colocalisation highlighted with yellow arrows. E. Inhibitor oligo nucleotides (D) were transfected into MCECs followed by UV-crosslinking, RNA interactome capture and Western blot analysis of the relative mRNA binding of Phf6 in the presence and absence of inhibitor. N=3, significance assessed by students t-test, error bars show SEM, significance shown to three sig. fig. F Inhibitor oligo nucleotides were transfected into MCECs, alongside siPhf6 as a control and rescue with wild-type Phf6 overexpression, for 24 hours followed by assessment of tubule formation. N=6, data shown as average ±SEM, significance assessed by one-way ANOVA, significance indicated.

### Ser199 phosphorylation promotes Phf6-RNA interactions in the cytoplasm

Utilising the specificity of RNA-bound Phf6 phosphorylation at S199, we used a classical mutagenesis approach to generate mutants of Phf6 mimicking either constitutively unphosphorylated (phospho-deficient) or constitutively phosphorylated (phospho-mimetic) Phf6 with S199A and S199E point mutations, respectively (Fig. 4C). AlphaFold analysis predicted the S199E mutation to result in a conformational shift, seemingly aligning the functional PHD domains and N-and C-terminal end peptides (Fig. 4D and Sup. Fig. 6C and D). After overexpressing wild type (WT), phospho-deficient (S199A) or phospho-mimetic (S199E) Phf6-myc in MCECs followed by RNA interactome capture and Western blotting for Phf6-myc, we found that S199E Phf6 exhibits significantly increased RNA binding affinity compared to WT and S199A Phf6 (Fig. 4E). Immunofluorescence microscopy of intracellular localisation and mRNA interactions confirmed increased mRNA binding in the S199E mutant, as well as an increased shift to the cytoplasm, additionally supported by nuclear-cytoplasmic fractionation and Western blot (Fig. 4F, Sup. Fig. 6E).

Phf6 canonically associates with transcriptional remodelling complexes such as the NuRD (Nucleosome Remodelling and Deacetylase complex) complex in the nucleus. Given that Ser199 phosphorylation induces a cytoplasmic shift in location and that Phf6 interacts with RNA primarily in the cytoplasm, we hypothesised that Ser199 phosphorylation results in a decrease in the interactions between Phf6 and its known nuclear binding partners (Fig. 4G). We tested if S199 phosphorylation affects the interaction of Phf6 with NuRD complex proteins by overexpressing wild type (WT), phospho-deficient (S199A) or phospho-mimetic (S199E) Phf6-myc followed by co-IP. S199 phospho-deficient Phf6 exhibited significantly increased interactions with HDAC (Histone deacetylase)1, HDAC2 and Chd4 (Chromodomain Helicase DNA-Binding Protein) compared to S199 phospho-mimetic Phf6 (Fig. 4H and I).

### Phf6 Ser199 phosphorylation is required to promote angiogenesis

We knocked down Phf6 in MCECs and examined the effects of rescue with either wild-type, phospho-deficient or phosphomimetic Phf6 on tube formation, proliferation and migration (Fig. 4 J, K and L). Whilst the loss of each of these pro-angiogenic traits could be rescued by wild-type (WT) or phospho-mimetic (S199E) Phf6, reintroducing phospho-deficient (S199A) Phf6 had no effect compared to Phf6 knock-down, suggesting that S199 phosphorylation plays an important role in driving a pro-angiogenic phenotype. Phf6 overexpression promotes angiogenic function also in ECs without Phf6 downregulation, but addition of the SRPK inhibitor abrogated the Phf6 driven increase in migration, proliferation and tube formation, suggesting that phosphorylation of Phf6 downstream of SRPK is required (Sup. Fig. 10 A-C). As expected, addition of an AKT activator could not rescue the SPRK inhibitor mediated effects, again indicating that SPRK acts downstream of AKT.

### Phf6 S199 phosphorylation promotes interaction with EMT and angiogenesis related RNAs

To investigate how Phf6 S199 phosphorylation alters the interaction between Phf6 and specific RNAs, we overexpressed wildtype (WT), phospho-deficient (S199A) or phospho-mimetic (S199E) Phf6 in MCECs followed by RNA immunoprecipitation and RNAseq analysis (Fig. 5A). In line with RNA interactome capture findings, this showed that phospho-mimetic Phf6 interacts with markedly more RNAs than wild-type or phospho-deficient Phf6 (816 vs. 172 or 136 RNAs, respectively (i.e. RNAs significantly enriched in Phf6 IPs compared to expression in matched inputs) (Fig. 5B and C). Enrichment analysis (MSigDB hallmark terms) showed a particular enrichment for RNAs involved in EMT, angiogenesis, apical junction and Notch signalling related pathways among the RNAs bound by phosphomimetic Phf6 (Fig. 5D). Interestingly, this strongly correlated with changes in expression of EMT and apical junction-related RNAs that occurred following VEGF-A stimulation (as well as following Phf6 knock-down).

Given this differential binding, we screened the enriched RNA ‘peaks’ identified in RIP-seq analysis for oligonucleotide motif enrichment, finding a clear enrichment of UAAU rich RNA sequences only in S199E-bound RNAs, suggesting that differential motif affinity may facilitate the differential binding, correlating with the change in Phf6 conformation, which demonstrated a closer interaction between UAAU repeats and the two PHD domains of Phf6 following phosphorylation (Fig. 5E-G). We found a particular increase in RNAs containing five or more UAAU motifs per kilobase, with these ‘hyper UAAU’ containing RNAs particularly enriched for the same angiogenesis and EMT-related functions (Fig. 5H and I). Together, this suggested that differential binding affinity for UAAU rich RNAs conferred by S199 phosphorylation may facilitate increased binding to angiogenesis and EMT-related RNAs.

### Phf6 differentially coordinates target RNA expression to promote angiogenesis

We overlapped the bound target RNAs of S199E Phf6 (all RNAs significantly enriched in Phf6 S199E IP vs input) with RNAs that showed opposing changes in expression after siRNA knockdown of Phf6 (i.e. from Fig. 3G) and upon overexpression of S199E Phf6 vs overexpression of S199A Phf6 (from analysis of RIPseq input samples, Sup. Fig. 9) (Fig. 6A).

This suggested that the RNA binding of Phf6 acts primarily to downregulate the expression of target RNAs as whilst 73 targets were downregulated by overexpression and upregulated by knockdown, only five S199E target RNAs were upregulated by overexpression and downregulated by Phf6 knock-down. Of these 73 S199E-Phf6 target RNAs which are downregulated by Phf6 there was primarily an enrichment of mesenchymal (EMT) along with apical junction and notch signalling related RNAs, many of which are known to be anti-angiogenic, including Col5a1 *(Collagen type-V alpha-1 chain)* and *Notch2*. In contrast only a rather small fraction of genes upregulated by S199E Phf6 and downregulated by siPhf6 were actually bound by S199E Phf6, indicating that induction of genes upon Phf6 phosphorylation might occur independent of RNA binding, for example due to altered Phf6 nuclear/chromatin regulating activity or secondary to changes in target RNA expression.

Nevertheless, among the five S199E-Phf6 target RNAs upregulated by Phf6 were *Mmp2 (Matrix Metalloproteinase-2)* and *Dag1 (Dystroglycan1),* which have known proangiogenic roles (Fig. 6A) (48,49). In addition, among the 280 genes upregulated, but not bound by S199E Phf6 were many angiogenesis promoting genes, i.e. *Angptl6*, *Bsg*, *Shc1*, *Tnfaip2*, *Vegfb*, *Rin2*, *Grn*, *Dab2*, *Rspo1* and *Sparc* whilst we found no angiogenesis, EMT or apical junction related genes among the 172 genes downregulated, but not bound by S199E Phf6. Following an alternative perhaps less stringent approach, we overlapped S199E Phf6 bound targets with the conserved siPHf6 regulated genes in HUVECS and MCECs and again identified Notch2 and Col5a1 as anti-angiogenic upregulated, as well Dag1 as pro-angiogenic downregulated direct Phf6 target.

This suggests a differential function of the binding of phosphorylated-Phf6 to target RNAs in order to facilitate a switch to a pro-angiogenic phenotype; on the one hand directly binding to and decreasing the expression of RNAs such as Col5a1 *(Collagen type-V alpha-1 chain)* and Notch2 which act as inhibitors of angiogenesis and EC proliferation and survival, whilst promoting the expression of RNAs such as *Mmp2* and *Dag1,* which can promote angiogenesis (48,50,51,52,53). In addition, the induction of pro-angiogenic RNAs by Phf6 may involve an additional, RBP independent mechanism that remains to be elucidated.

We chose three RNAs, which showed differential binding patterns with distinct functions in angiogenesis and EMT for further validation and characterisation. The relative binding of Phf6 to *Notch2* was significantly enriched for both wild-type (WT) and phospho-mimetic (S199E) Phf6 IPs, whereas binding to *Col5a1* was significantly enriched only in S199E IPs and interaction with *Mmp2* RNA was elevated in all conditions (though not significant) as assessed by RIP followed by qPCR (RIP-qPCR) (Fig. 6B). RIP-qPCR of the binding of endogenous Phf6 to these targets showed a significant interaction with *Notch2* and *Col5a1* after VEGF-A stimulation, whereas *Mmp2* binding was significant whether or not VEGF-A was present and was constitutive in MCECs (Fig. 6C). Interestingly, AKT activation promoted binding to these targets, which was inhibited by SRPK inhibition (Sup. Fig. 10D). In HUVECs, we found that VEGF-A drives a significant increase in binding to *NOTCH2* and *COL5A1*, and a significant decrease in binding to *MMP2,* with AKT OE in HUVECs resulting in a significant increase in binding to *COL5A1* and a moderate increase (not significant) to *NOTCH2* and *MMP2* (Fig. 6D, Sup. Fig. 7C).

We investigated the effects of Phf6, and S199 phosphorylation, on the expression of *Notch2*, *Col5a1* and *Mmp2*, finding that knockdown of Phf6 resulted in a significant increase in expression of anti-angiogenic *Notch2* and *Col5a1* and a decrease in pro-angiogenic *Mmp2* expression, which in all cases was rescued only by wildtype or phosphomimetic Phf6, but not by the phosphodeficient form, suggesting that S199 phosphorylation may play a direct role in coordinating the expression of these target RNAs (Fig. 6E). This again suggests that S199 phosphorylation simultaneously acts as a switch to promote expression of pro-angiogenic targets whilst decreasing the expression of anti-angiogenic targets.

### Phf6 driven decrease in Col5a1 and Notch2 promotes tube formation

To analyse whether the increase in Col5a1 and Notch2 promote anti-angiogenic features, when Phf6 is downregulated, we performed a tube formation assay following knock-down of Phf6 as well as Notch2 or Col5a1 knockdown. We found a significant rescue (i.e. increase) in tube formation with combined Col5a1 and Notch2 knockdown during Phf6 knock-down (Fig. 6F). This suggested that Phf6-mediated changes in target RNA expression play a role in coordinating the proangiogenic phenotype associated with Phf6 activation.

### Differential Phf6 binding coordinates changes in target RNA stability

Given the direct impact of Phf6 on the expression of *Notch2, Col5a1* and *Mmp2*, we hypothesised that this may be due to differential effects of Phf6 on the stability of these target RNAs. Using an Actinomycin D (an inhibitor of *de novo* RNA transcription) time-course followed by calculation of the respective decay rate of each RNA, we found that S199 phosphorylation of Phf6 specifically increases the decay rate of *Notch2* and *Col5a1*, whilst promoting *Mmp2* stability (Fig. 7A). As we found a link between Phf6 S199 phosphorylation and the presence of UAAU motifs within target RNAs, we visualised the occurrence of UAAU motifs in *Notch2, Col5a1* and *Mmp2,* finding that *Notch2* and *Col5a1* had a distinctly high number of UAAU motifs in their 3’UTR region (Fig. 7B). As *Notch2* and *Col5a1* were both destabilised by our S199 phosphomimetic, we hypothesised that S199 phosphorylation may drive a differential interaction of Phf6 with the 3’UTRs of these RNAs to promote their decay. To test this, we cloned the 3’UTR of *Notch2* into a luciferase reporter assay, which was transfected into MCECs alongside wild-type, phosphodeficient or phosphomimetic Phf6 constructs (Fig. 7C). This showed that whilst Phf6 knockdown significantly increased reporter stability, this was rescued specifically by wild-type or phosphomimetic Phf6, but not by phosphodeficient Phf6. This suggests that upon S199 phosphorylation Phf6 differentially binds to UAUU motifs within the 3’UTR of target mRNAs to decrease their stability.

Because our data suggested that Phf6 phosphorylation may confer differential binding affinity for specific RNA motifs (e.g. higher affinity for UAAU rich RNAs upon S199 phosphorylation), we aimed to utilise this information to test the principle of competitive oligonucleotide inhibitors to regulate the RNA binding of Phf6. We designed two oligonucleotides, Oligo1 based on the UAAU rich motifs primarily bound by phosphomimetic Phf6 and Oligo2 based on the motifs primarily bound by wild-type and phosphodeficient Phf6, each harbouring a biotin tag for capture and visualisation (Fig. 7D).

Immunofluorescence microscopy confirmed a co-localisation between the oligos and Phf6, and suggested an increased interaction with Oligo1 upon VEGF-A stimulation, correlating with Phf6 phosphorylation, while Oligo2 binding of Phf6 did not respond to VEGF-A (Fig. 7D). Co-immunoprecipitation of either Phf6 or biotinylated oligos confirmed a stronger binding affinity between Phf6 with Oligo1 than Oligo2, a preferential interaction between wild-type and phosphomimetic Phf6 with Oligo1 and a decrease in affinity for phosphomimetic Phf6 with Oligo2 (Sup. Fig. 11 A and B).

We next tested whether the Oligos could compete with endogenous RNA for mRNA binding. Indeed, transfection of Oligo1 significantly impaired the global RNA interactions of phosphomimetic Phf6, which was unaffected by Oligo2 transfection, suggesting that the increased interactions of Phf6 with RNA upon S199 phosphorylation are at least in part driven by the increased binding affinity for UAAU rich regions (Fig. 7E).

Finally, we tested if the transfection of the competitive Oligos into MCECs would inhibit tube formation comparably to siRNA knockdown of Phf6, finding that only transfection with oligo 1 (i.e. inhibiting phosphorylation driven increases in Phf6-RNA interactions) inhibited EC tube formation in a similar manner to siPhf6 knockdown (Fig. 7F).

In summary, VEGF-A stimulation results in highly dynamic changes in in RBP phosphorylation at the global level. Specifically, phosphorylation of Phf6 at Ser199 promotes its interaction with the 3’UTR of anti-angiogenic RNAs, decreasing their stability and expression, whilst at the same time promoting the expression of proangiogenic RNAs, in part independent of its RNA binding function.

## DISCUSSION

RBP phosphorylation plays a key role in determining changes in RBP function, such as altering RBP-RNA localisation, binding affinity for subsets of target RNAs, and coordinating changes in RNA translation, splicing and stability. Recent approaches have revealed that global changes in the phosphorylation of the RNA-bound proteome have regulatory potential (5,8,54). We therefore hypothesised that dynamic changes of RNA-bound proteins downstream of VEGF-A stimulation may act to coordinate angiogenesis at the post-transcriptional level in ECs. By combining RNA interactome capture with simultaneous proteomic and phosphoproteomic analysis, we set out to quantify the differential phosphorylation of RBPs occurring in response to acute and chronic VEGF-A stimulation in the global mRNA-bound proteome.

VEGF-A signals predominantly through VEGFR2, initiating receptor autophosphorylation, which activates multiple parallel signalling cascades, including the MAPK/ERK, PI3K-AKT, PLCγ, FAK/Src and STAT3 pathways (55). More than 240 proteins and their post-translational modification-dependent interactions have now been catalogued across the VEGFA/VEGFR2 network, demonstrating the importance of PTMs in coordinating angiogenic responses (56). We found that stimulation with either 15 minutes or 24 hours of VEGF-A resulted in temporally specific changes in both RBP phosphorylation and total RNA-binding patterns, reflecting two functionally and mechanistically distinct phases of the endothelial response to VEGF-A. The 15-minute timepoint captures rapid post-translational remodelling driven by activation of intracellular kinases such as Erk1/2, FAK and PI3K-AKT downstream of VEGFR2 phosphorylation (57,58). In contrast, the 24-hour timepoint reflects a later phase of transcriptional and translational change required for angiogenic progression, executed in large part through a defined set of angiogenic transcription factors such as ETS1, ERG, MEF2C and FOXO1 (59,60,61).

Accordingly, we identified a number of RBPs that were differentially phosphorylated exclusively at either 15 minutes or 24 hours post-VEGF-A, whereas other differentially phosphorylated RBPs remained altered at both timepoints, suggesting a more prolonged role in angiogenic remodelling. The predicted upstream kinases of differentially phosphorylated RBPs also reflected known downstream effector kinases of VEGF-A (e.g. MTOR, AKT, PAK1, ERK1, JNK1), providing insights into how they may coordinate differential phosphorylation of the RNA interactome at these distinct timepoints. This positions RBP phosphorylation as a regulatory layer that operates in parallel with transcriptional reprogramming. We found that the differential phosphorylation of certain RBPs directly correlated with binding occupancy on RNA (i.e. more or less phosphorylation directly correlated with more or less binding to RNA at the global level), whereas others showed no significant differences in their abundance in global mRBP isolates (i.e. the RBP was differentially phosphorylated, but the global binding activity did not change). This suggests alternative regulatory impacts of these phosphorylation events, as phosphorylation may result in changes in the total binding of the RBP to target RNAs or regulate other functions such as interactions with specific subsets of target RNAs, interactions with other proteins or changes in metabolic activity to coordinate the endothelial response to VEGF-A and promote angiogenesis.

We detected phosphosites within almost two-thirds of the mRNA-bound proteome covering almost all aspects of RNA metabolism including the regulation of RNA translation, splicing, stability and localisation. A large number of differentially phosphorylated sites following VEGF-A stimulation were detected within two RNA-binding proteins, Serine and Arginine Repetitive Matrix (Srrm) 1 and Srrm2, splicing factors known to be highly regulated by phosphorylation to coordinate cellular responses to stress and signalling cues (62,63,64). In contrast, the majority of RBPs contained a small number of phosphosites, suggesting the capacity for more specific regulation.

We found only one phosphorylation site within Phf6 (PHD Finger Protein 6) that showed the most pronounced increase in phosphorylation of all RBPs after both 15 minutes and 24 hours of VEGF-A stimulation. Phf6 is a highly conserved PHD finger-containing protein best known for its roles in chromatin regulation, transcriptional control and ribosomal biogenesis, with loss-of-function mutations linked to multiple diseases, particularly as a driver of various forms of leukaemia (21,65). More recently, Phf6 has also emerged in multiple RNA interactome studies, as well as other methods of RBP identification such as phase-separation and centrifugation-based methodologies, demonstrating a direct association with mRNA (e.g. 25,26,27,28). Whilst the detailed role of Phf6 as an mRNA-binding protein remains to be fully characterised, eCLIP-seq (Enhanced Crosslinking and Immunoprecipitation Sequencing) data has recently revealed that Phf6 directly interacts with the 3’UTR, 5’UTR and coding sequence of protein-coding RNAs, as well as rRNAs and other non-coding RNAs (66).

Our data identified Ser199 as the exclusive phosphorylation site within mRNA-bound Phf6 upon VEGF-A stimulation in ECs, suggesting that Ser199 plays an important role in promoting the interaction between Phf6 and mRNA. Our data suggest that SRPK kinases act as the likely upstream drivers of Ser199 phosphorylation, downstream of AKT. SRPK kinases, located in both the nucleus and cytoplasm, are known to be key regulators of RNA metabolism through phosphorylation of RS (arginine/serine) dipeptides within target proteins (67). The best-characterised role of SRPK kinases is in phosphorylating serine– arginine-rich splicing factors (SRSFs) to coordinate RNA splicing patterns, which is known to be pro-angiogenic by facilitating alternative VEGF splicing downstream of SRSF1 (68,69,70). Together with this, our findings suggest that SRPK kinases may have more diverse effects on RBP phosphorylation in promoting angiogenesis than currently characterised, which merits further investigation.

We found that Phf6 phosphorylation at Ser199 correlated with both a change in Phf6 localisation and a dramatic change in the RNAs with which Phf6 interacts. Phf6 has been shown to locate to the nucleolus, nucleus and cytoplasm (66), and an RNA-dependent effect on Phf6 localisation has previously been reported, with RNA interaction shown to mediate a shift in Phf6 localisation to the nucleolus (71). Phf6 canonically functions as a chromatin adaptor protein that interacts with several transcriptional regulatory complexes, directly binding the NuRD complex to mediate transcriptional regulation at target genes (22,65).

Interestingly, we revealed that Ser199 phosphorylation reduces the interaction with various NuRD complex components, correlating with an increase in cytoplasmic Phf6 localisation. It could therefore be that a change in RNA binding, possibly driven by a phosphorylation-induced conformational change that preferentially confers a pro-RNA-binding conformation, facilitates a change in Phf6 localisation; or, conversely, that a change in Phf6 localisation, for example due to decreased interaction with nuclear complexes, facilitates a change in RNA-binding patterns; or that the two effects act synergistically.

Whilst a direct role for Phf6 in VEGF-A-driven angiogenesis has not previously been investigated, several studies in other models have reported roles for Phf6 that may align with our findings that Phf6 promotes angiogenesis and EC proliferation. For example, Phf6 can physically interact with HIF-1α and HIF-2α to induce downstream HIF signalling events and promote cancer progression, while downregulation of Phf6 inhibits proliferation and migration in a hepatocellular carcinoma model (72,73). Several studies have also revealed that phosphorylation may play an important role in regulating Phf6 function: phosphorylation at Ser145, Ser154 and Ser155 occurs during mitosis and cell cycle progression (21,74), and phosphorylation of five serine residues, including Ser199, has been reported in Phf6 in response to DNA damage, with Ser199 phosphorylation inhibited by caffeine treatment (23).

We found that loss of Phf6 increased the expression of target RNAs involved in EMT and apical junction organisation, and decreased those involved in promotion of angiogenesis, KRAS, mTORC1 and mitotic signalling, opposing the transcriptomic changes induced by VEGF-A. This suggests that VEGF-A-driven Phf6 phosphorylation and increased RNA binding act as a switch, decreasing anti-angiogenic RNAs and EMT promoting RNAs by fostering their degradation. S199 phosphorylation of Phf6 on the other hand promotes pro-angiogenic RNA expression, in large parts independent of its RNA binding function.

The RNAs that are repressed by Phf6 include a large number of pro-mesenchymal EMT-annotated RNAs, many of which overlap with EMT/endothelial-mesenchymal transition (EndoMT) annotations and contribute to ECM remodelling and the inhibition of EC migration and invasion (75,76). *Col5a1*, encoding a fibrillar collagen, was increased following Phf6 loss and was exclusively bound by Phf6 upon S199 phosphorylation, consistent with reports that type V collagen, and Col5a1-derived fragments, in the endothelial basement membrane restrain pathological angiogenesis (50,51). Phf6 may therefore bind to and decrease matrix-related gene expression to permit EC migration and sprouting upon its phosphorylation at S199. Likewise, phosphorylated Phf6 binds to and decreases *Notch2*, a reported inhibitor of AKT signalling, EC survival and proliferation (52,77). Since Phf6 Ser199 phosphorylation occurs downstream of AKT, Notch2 repression may reinforce the same signalling axis that drives Phf6 phosphorylation.

Conversely, the downregulation of KRAS, mTORC1 and mitotic signalling RNAs upon Phf6 loss is consistent with a role for Phf6 in sustaining proliferative, growth-factor-responsive programmes, as VEGFR2 activation drives both PI3K/AKT/mTORC1 and RAS/ERK signalling to coordinate EC proliferation and vessel growth (78,79). *Mmp2* and *Dag1*, target RNAs of phosphorylated Phf6, were decreased upon Phf6 knockdown. Mmp2 is a matrix-remodelling protease that promotes neovascularisation, whereas Dag1 can stabilise EC sprouts and promote migration (48,49,80).

Intriguingly, however, most of the upregulated pro-angiogenic mRNAs were not directly bound by S199-Phf6, implying either indirect effects of RNA bound Phf6 or direct transcriptional effects, but further work is needed to decipher the exact mechanisms. We postulate that this may be due to its altered function as a transcriptional regulator, for example due reduced suppression via the NuRD complex. Together, these data position Phf6 as a key regulator of VEGF-A-driven expression programmes, in which phosphorylation couples the repression of anti-angiogenic RNAs to the maintenance of pro-angiogenic ones.

Mechanistically, we found that phosphorylation of Ser199 promoted distinct changes in RNA-binding patterns, facilitating interaction, potentially with specific motifs, within subsets of target RNAs, directly correlating with changes in their stability and expression. In particular, we propose that Ser199 phosphorylation drives differential binding of Phf6 to UAAU motifs within the 3’UTR of specific target transcripts which are primarily inhibitory to angiogenesis (e.g. RNAs with pro-mesenchymal functions) in order to decrease their stability, whilst synergistically increasing the expression of proangiogenic RNAs. This provides a putative mechanism through which phosphorylation-dependent changes in Phf6-RNA interactions act as a switch in order to into differentially orchestrate changes in target mRNA stability and expression to drive the proangiogenic phenotype associated with S199 phosphorylation. This would be consistent with a broader precedent for short AU-rich 3’UTR elements acting as binding sites for RBPs that directly govern transcript stability: AU-rich elements (AREs) are bound by ARE-binding proteins that accelerate, or in some contexts stabilise, degradation of their target transcripts (81,82).

The use of RNA-based ’decoy’ inhibitors, RNA oligonucleotides designed to act as competitive inhibitors of RBP-RNA interactions, is emerging as an effective way to modulate RBP function (83). Accordingly, we showed that transfection of an RNA decoy could effectively inhibit Phf6-RNA interactions and angiogenic features, suggesting the possibility of specifically inhibiting the RNA-binding function of Phf6.

Together, we show that stimulation of endothelial cells with VEGF-A results in widespread and dynamic changes in the phosphorylation of the RNA-bound proteome to coordinate pro-angiogenic processes at the post-transcriptional level. We highlight the role of Ser199 phosphorylation within RNA-bound Phf6 as an important pro-angiogenic switch between its canonical function as a transcriptional regulator with the nucleus to an mRNA binding protein in the cytoplasm where it coordinates the inhibition of anti-angiogenic RNAs. Whilst most studies of Phf6 have focused on its role in transcriptional complexes and ribosome biogenesis, further investigation into its context-dependent RNA-binding patterns, and phosphorylation, may add previously unstudied insight into the role of Phf6 in various cellular and disease models. We suggest that our global phosphoproteomic data on VEGF-A-driven changes in RBP phosphorylation may act as a platform for future investigations into the importance of changes in RNA-binding patterns in coordinating angiogenesis and other endothelial responses.

## ACKNOWLEDGEMENTS

The authors gratefully acknowledge the data storage service SDS@hd supported by the Ministry of Science, Research and the Arts Baden-Württemberg (MWK) and the German Research Foundation (DFG) through grant INST 35/1314-1 FUGG and INST35/1503-1 FUGG.

## AUTHOR CONTRIBUTIONS

RW and JH: conceptualisation, methodology, formal analysis, visualisation, writing-original draft and final draft

JH: Funding and Supervision

LI, AT, KB, SJ, JS, FS: methodology, formal analysis, visualisation RO, GD, AH and JK: methodology, writing-review and editing

## SUPPLEMENTARY DATA

Supplementary Data are available at NAR online.

