## Supplemental Figures for "Global mapping of VEGF-A driven changes in the RNA bound phospho-proteome reveals a molecular switch to coordinate angiogenesis in endothelial cells"

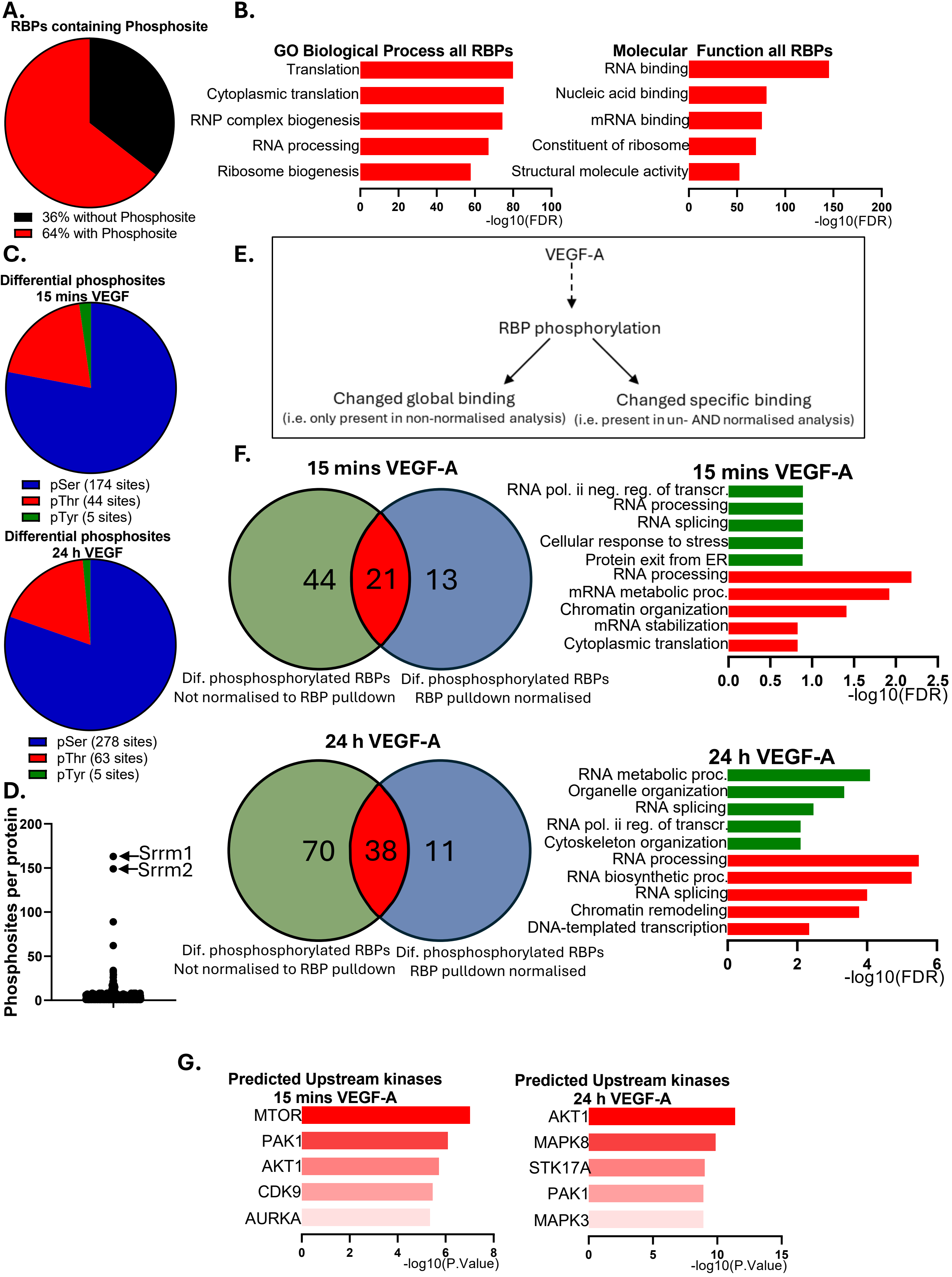

A.
RBPs containing Phosphosite
36% without Phosphosite
64% with Phosphosite
B.
GO Biological Process all RBPs
Translation
Cytoplasmic translation
RNP complex biogenesis
RNA processing
Ribosome biogenesis
0 20 40 60 80 100
-log10(FDR)
Molecular Function all RBPs
RNA binding
Nucleic acid binding
mRNA binding
Constituent of ribosome
Structural molecule activity
0 50 100 150 200
-log10(FDR)
C.
Differential phosphosites 15 mins VEGF
pSer (174 sites)
pThr (44 sites)
pTyr (5 sites)
Differential phosphosites 24 h VEGF
pSer (278 sites)
pThr (63 sites)
pTyr (5 sites)
D.
Phosphosites per protein
0 50 100 150 200
Srrm1
Srrm2
E.
VEGF-A
RBP phosphorylation
Changed global binding
(i.e. only present in non-normalised analysis)
Changed specific binding
(i.e. present in un- AND normalised analysis)
F.
15 mins VEGF-A
44 21 13
Dif. phosphosphorylated RBPs Not normalised to RBP pulldown
Dif. phosphosphorylated RBPs RBP pulldown normalised
15 mins VEGF-A
RNA pol. ii neg. reg. of transcr.
RNA processing
RNA splicing
Cellular response to stress
Protein exit from ER
RNA processing
mRNA metabolic proc.
Chromatin organization
mRNA stabilization
Cytoplasmic translation
0.0 0.5 1.0 1.5 2.0 2.5
-log10(FDR)
24 h VEGF-A
70 38 11
Dif. phosphosphorylated RBPs Not normalised to RBP pulldown
Dif. phosphosphorylated RBPs RBP pulldown normalised
24 h VEGF-A
RNA metabolic proc.
Organelle organization
RNA splicing
RNA pol. ii reg. of transcr.
Cytoskeleton organization
RNA processing
RNA biosynthetic proc.
RNA splicing
Chromatin remodeling
DNA-templated transcription
0 2 4 6
-log10(FDR)
G.
Predicted Upstream kinases 15 mins VEGF-A
MTOR
PAK1
AKT1
CDK9
AURKA
0 2 4 6 8
-log10(P.Value)
Predicted Upstream kinases 24 h VEGF-A
AKT1
MAPK8
STK17A
PAK1
MAPK3
0 5 10 15
-log10(P.Value)

**Supplemental Figure 1.** Extended proteomic analysis related to Figure 1. A. Proportion of RBPs identified containing one or more phosphorylation sites. B. Enriched GO biological process and molecular function terms of RBPs identified in proteomic analysis. C. Proportion of differentially phosphorylated Ser, Thr and Tyr residues within RBPs after 15 minutes or 24 hours of VEGF-A stimulation. D. The number of individual phosphosites within each phosphorylated RBP. E. Overview of potential effects of phosphorylation on RNA binding patterns. F. Proportion of differentially phosphorylated RBPs identified with and without normalisation to changes in the corresponding input ‚RBPomes', with corresponding analysis of enriched GO biological processes (significant differential phosphorylation sites occurring in unnormalised (green) and normalised (red) analysis). G. Predicted upstream kinases of differential changes in RBP phosphorylation after 15 minutes or 24 hours of VEGF-A stimulation, after screening differential phospho-sites against the PhosD database.

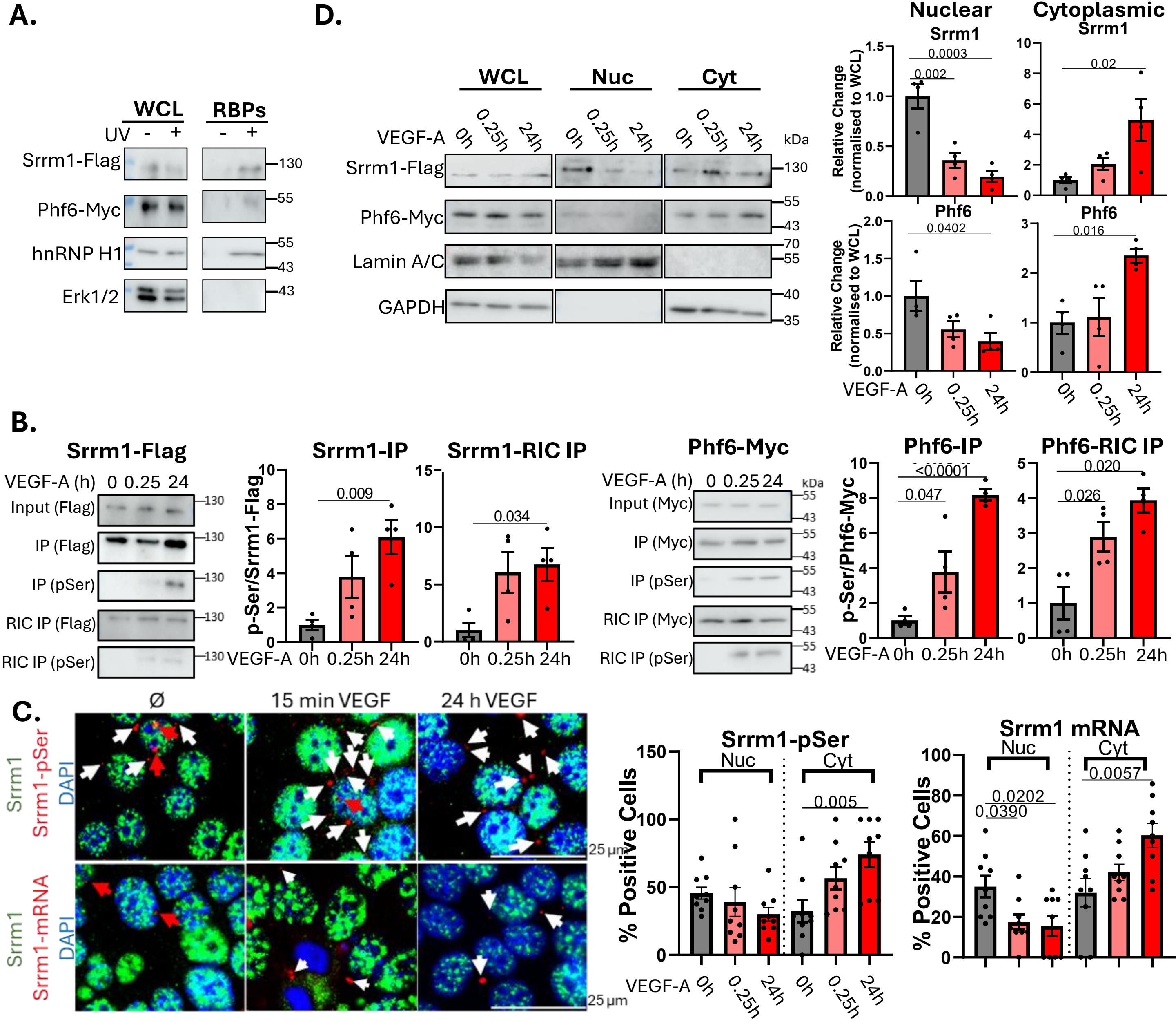
A.
WCL
RBPs
UV - + - +
Srrm1-Flag
Phf6-Myc
hnRNP H1
Erk1/2
130
55
55
43
43
D.
WCL
Nuc
Cyt
VEGF-A 0h 0.25h 24h 0h 0.25h 24h 0h 0.25h 24h
kDa
Srrm1-Flag
Phf6-Myc
Lamin A/C
GAPDH
130
55
43
70
55
40
35
Nuclear
Cytoplasmic
Srrm1
Phf6
Relative Change (normalised to WCL)
0.0003
0.002
0.02
0.0402
0.016
VEGF-A 0h 0.25h 24h
B.
Srrm1-Flag
VEGF-A (h) 0 0.25 24
Input (Flag)
IP (Flag)
IP (pSer)
RIC IP (Flag)
RIC IP (pSer)
Srrm1-IP
Srrm1-RIC IP
p-Ser/Srrm1-Flag
0.009
0.034
VEGF-A 0h 0.25h 24h
Phf6-Myc
Input (Myc)
IP (Myc)
IP (pSer)
RIC IP (Myc)
RIC IP (pSer)
Phf6-IP
Phf6-RIC IP
p-Ser/Phf6-Myc
<0.0001
0.047
0.020
0.026
C.
Ø
15 min VEGF
24 h VEGF
Srrm1
Srrm1-pSer
DAPI
Srrm1-mRNA
25 µm
Srrm1-pSer
Nuc
Cyt
% Positive Cells
0.005
Srrm1 mRNA
0.0202
0.0390
0.0057
VEGF-A 0h 0.25h 24h 0h 0.25h 24h

**Supplemental Figure 2.** Extended validation and characterisation of Srrm1 and Phf6. A. Validation of Srrm1 and Phf6 as RBPs. RNA interactome capture was performed in MCECs in the presence and absence of UV-crosslinking, Western blot performed on RBP eluates. hnRNP H1 was included as a postive control and ERK 1/2 as a negative control. Representative blot from three replicates. B. Effects of VEGF-A stimulation on the nuclear/cytoplasmic localisation of Phf6 and Srrm1. Srrm1 (flag-tagged) or Phf6 (Myc tagged) were overexpressed in MCECs in the presence of either 15 minutes or 24 hours of VEGF-A stimulation, followed by nuclear-cytoplasmic fractionation and Western blot analysis (GAPDH and Lamin A/C were included as cytoplasmic/nuclear markers respectively). n=4, significance assessed by one-way ANOVA, significance indicated. C. Visualisation of VEGF-A driven changes in the phosphorylation and mRNA interactions of Srrm1. A proximity ligation assay (PLA) in MCECs stimulated with either 15 minutes or 24 hours of VEGF-A stimulation (10 ng/ml) using an antibody recognising endogenous Srrm1, along with either a p-Ser probe or biotinylated oligo d(T) probe followed by anti-biotin antibody, followed my immunofluorescence microscopy, was to visualise changes in the phosphorylation and RNA interactions of Srrm1 in MCECs following 15 minutes or 24 hours of VEGF-A stimulation. PLA signal (phosphorylated Srrm1 or Srrm1–mRNA interactions) is shown in red, total Srrm1 shown in green (DAPI in blue). White arrows show examples of cytoplasmic PLA signal and red arrows show examples of nuclear PLA signal. Quantifications show percentage of cells per image positive for nuclear and/or cytoplasmic PLA signal (n=9, significance assessed by one-way ANOVA, significance indicated). D. Validation of changes in phosphorylation. Srrm1 (flag-tagged) or Phf6 (Myc tagged) were overexpressed in MCECs followed by 15 minutes or 24 hours of VEGF-A stimulation, and immunoprecipitated from whole cell lysate or isolated RBPs (RIC isolates). Western blots were used to detect the relative amounts of serine phosphorylation (n=4, significance assessed by one-way ANOVA, significance indicated.

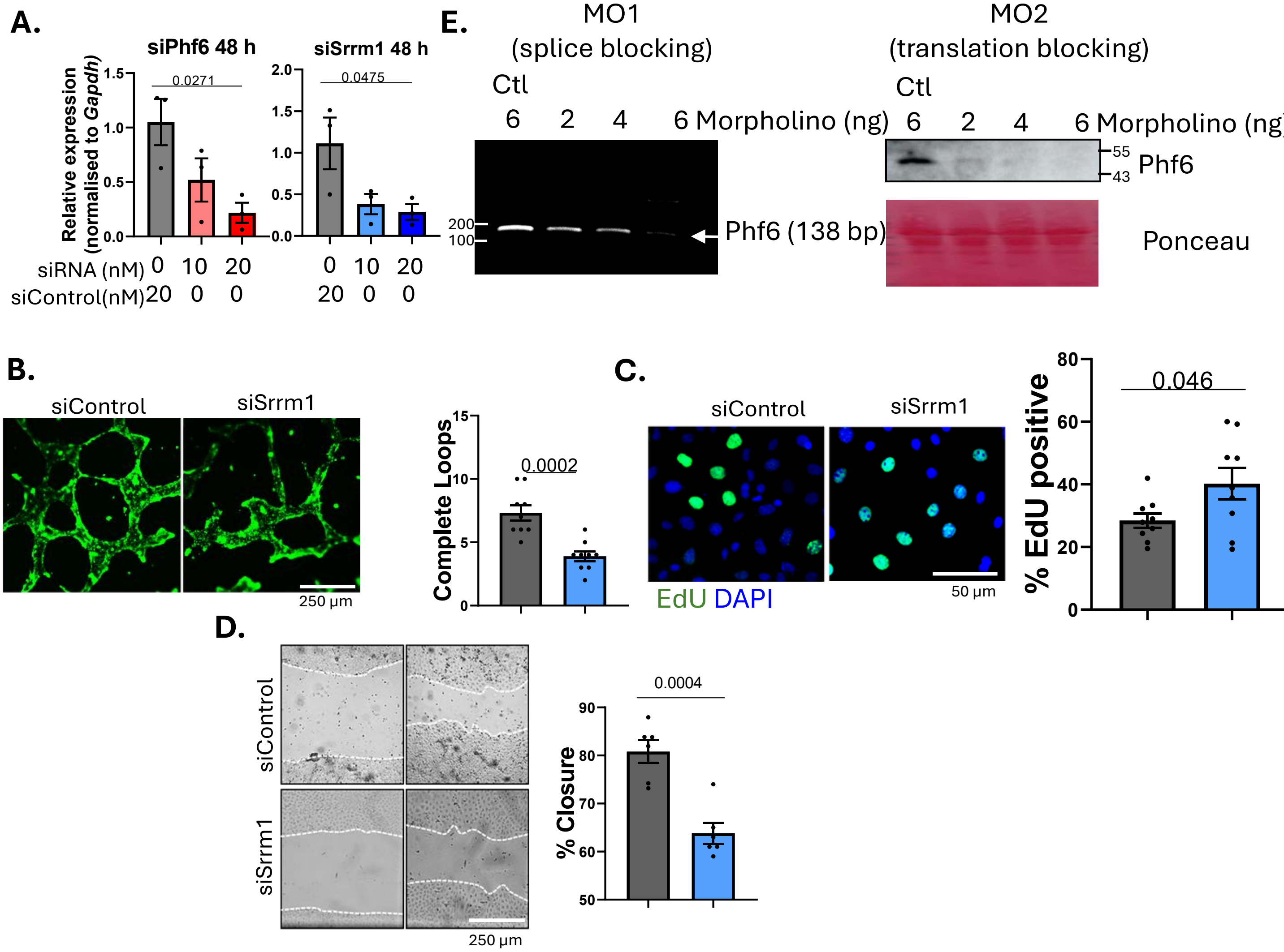


**Supplemental Figure 3.** A. Validation of Srrm1 and Phf6 siRNA. MCECs were transfected with siRNA against either Srrm1 or Phf6 for 48 hours (alongside scrambled control) followed by qPCR analysis of relative expression (normalised to *Gapdh)*. n=3, significance assessed by one-way ANOVA, significance indicated. B. Effects of Srrm1 knockdown (siRNA, 48 hours, ± 10 ng/ml VEGF-A stimulation for 24 h) on tube formation in MCECs plated in Matrigel for 24 hours. Tubes were fluorescently labelled with BCECF AM Ester prior to imaging. Representative images and quantifications of the number of complete loops per visible field, n=3 triplicates/condition, data shown as average ±SEM significance assessed by t-test, significance indicated. C. Effects of Srrm1 knockdown (siRNA, 48 hours, ± 10 ng/ml VEGF-A stimulation for 24 h) on EdU incorporation (proliferation) in MCECs. Representative images and quantifications of EdU incorporation, n=3 triplicates/condition, quantified as percentage EdU positive cells per image data shown as average ±SEM significance assessed by t-test, significance indicated. D. Effects of Srrm1 knockdown (siRNA, 48 hours, ± 10 ng/ml VEGF-A stimulation for 24 h) on MCEC migration as assessed by scratch wound assay. Representative images and quantification of the percentage closure 24 hours post-scratch, n=3 triplicates/condition. Data shown as average ±SEM significance assessed by t-test, significance indicated. E. Validation of loss of Phf6 in zebrafish using a morpholino based approach. Two independent morpholinos (MO1; splice blocking or MO2; translation blocking) were used and injected into the yolk sack of one-cell stage embryos. Phf6 expression was assessed 48 hpf by either PCR (MO1) or Western blot (MO2).

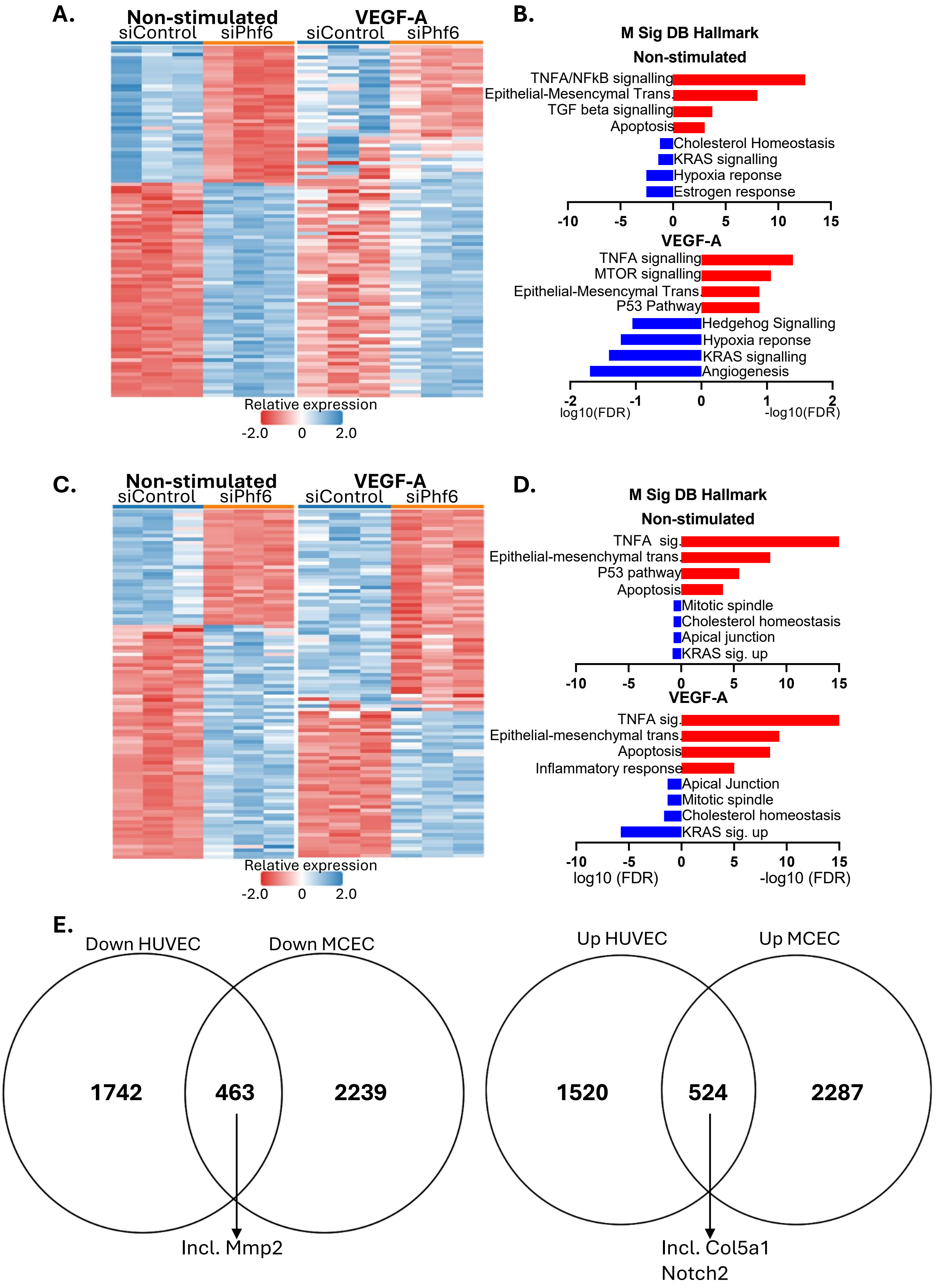
A.
Non-stimulated
siControl siPhf6
VEGF-A
siControl siPhf6
Relative expression
-2.0 0 2.0
B.
M Sig DB Hallmark
Non-stimulated
TNFA/NFkB signalling
Epithelial-Mesencymal Trans.
TGF beta signalling
Apoptosis
Cholesterol Homeostasis
KRAS signalling
Hypoxia reponse
Estrogen response
-10 -5 0 5 10 15
VEGF-A
TNFA signalling
MTOR signalling
Epithelial-Mesencymal Trans.
P53 Pathway
Hedgehog Signalling
Hypoxia reponse
KRAS signalling
Angiogenesis
-2 -1 0 1 2
log10(FDR) -log10(FDR)
C.
Non-stimulated
siControl siPhf6
VEGF-A
siControl siPhf6
Relative expression
-2.0 0 2.0
D.
M Sig DB Hallmark
Non-stimulated
TNFA sig.
Epithelial-mesenchymal trans.
P53 pathway
Apoptosis
Mitotic spindle
Cholesterol homeostasis
Apical junction
KRAS sig. up
-10 -5 0 5 10 15
VEGF-A
TNFA sig.
Epithelial-mesenchymal trans.
Apoptosis
Inflammatory response
Apical Junction
Mitotic spindle
Cholesterol homeostasis
KRAS sig. up
-10 -5 0 5 10 15
log10 (FDR) -log10 (FDR)
E.
Down HUVEC
Down MCEC
1742
463
2239
Incl. Mmp2
Up HUVEC
Up MCEC
1520
524
2287
Incl. Col5a1
Notch2

**Supplemental Figure 4.** Extended RNA sequencing analysis related to Figure 3. RNA sequencing analysis of the effects of Phf6 knockdown in MCECs and HUVECs. A. Heatmaps showing differential RNA expression patterns in MCECs following siRNA knockdown of Phf6 in the presence and absence of VEGF-A stimulation (48 hours of Phf6 knockdown (siPhf6), 24 hours of VEGF-A stimulation). B. Enriched M Sig DB Hallmark terms of differentially expressed RNAs ($p<0.05$, Log2 FC<0.5, red up- and blue downregulated by siPhf6) following siRNA knockdown of Phf6 in the presence and absence of VEGF-A stimulation (48 hours of Phf6 knockdown (siPhf6), 24 hours of VEGF-A stimulation) in MCECs. C. Heatmaps showing differential RNA expression patterns in HUVECs following siRNA knockdown of Phf6 in the presence and absence of VEGF-A stimulation (48 hours of Phf6 knockdown (siPhf6), 24 hours of VEGF-A stimulation). D. Heatmaps showing differential RNA expression patterns in HUVECs following siRNA knockdown of Phf6 in the presence and absence of VEGF-A stimulation (48 hours of Phf6 knockdown (siPhf6), 24 hours of VEGF-A stimulation). D. Enriched M Sig DB Hallmark terms of differentially expressed RNAs ($p<0.05$, Log2 FC<0.5, red up- and blue downregulated by siPhf6) following siRNA knockdown of Phf6 in the presence and absence of VEGF-A stimulation (48 hours of Phf6 knockdown (siPhf6), 24 hours of VEGF-A stimulation) in HUVECs. E. Overlap of all differentially expressed RNAs in HUVECs and MCECs following 48 hours of Phf6 knockdown (siPhf6)and 24 hours of VEGF-A stimulation ($P<0.05$).

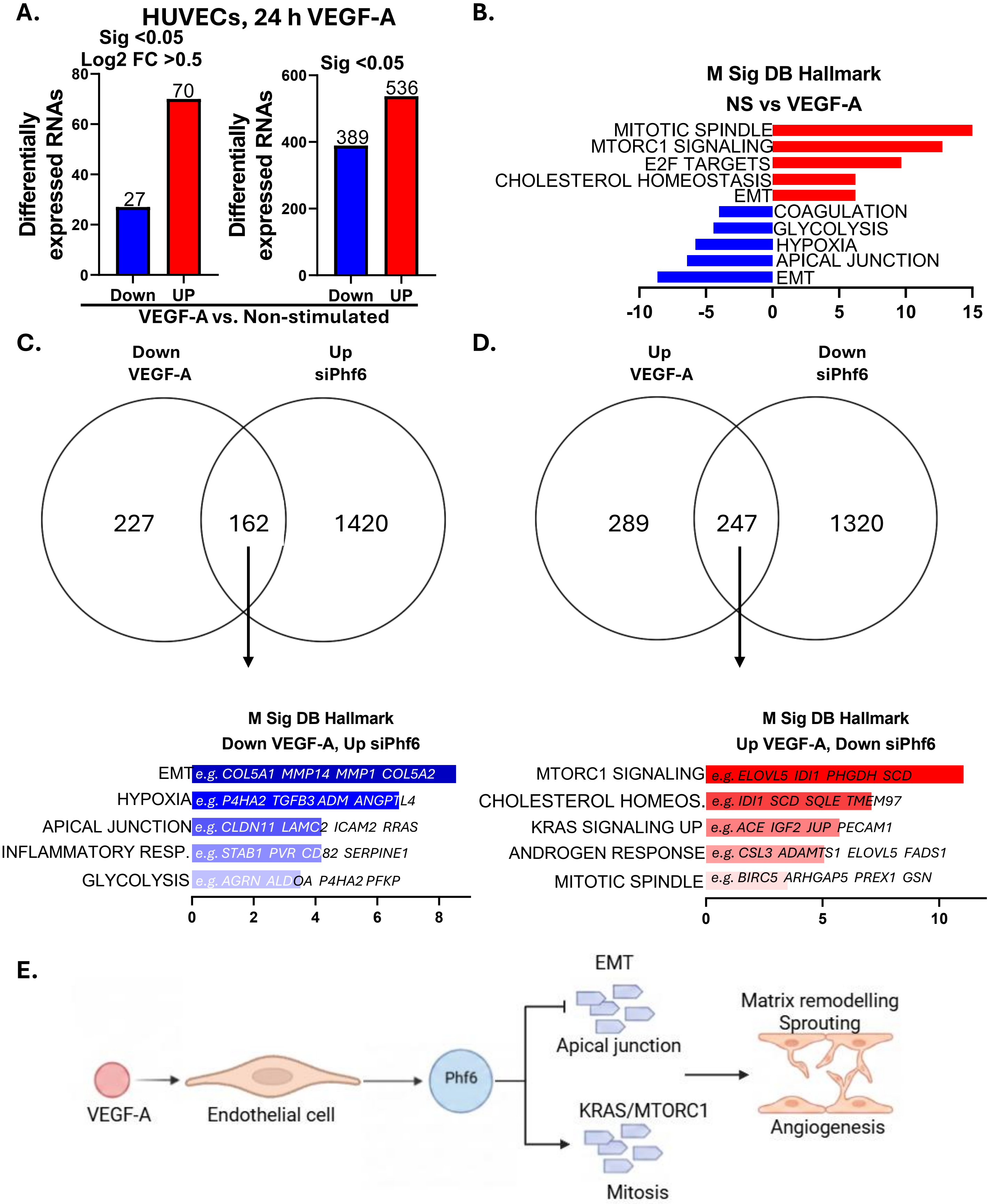


**Supplemental Figure 5. The role of Phf6 in mediating VEGF-A driven changes in RNA expression**. A. The number of up- and down-regulated RNAs following VEGF-A stimulation (24 h, 10 ng/ml) in HUVECs. B. Enriched M Sig DB Hallmark terms of differentially expressed RNAs (p<0.05) in HUVECs after VEGF-A stimulation. C and D. Overlap of differentially expressed RNAs (P<0.05) that were either downregulated by VEGF-A and upregulated by siPhf6 (C) or upregulated by VEGF-A and downregulated by siPhf6 (D) with corresponding enriched M Sig DB Hallmark terms. E. Schematic representation of the potential mechanism by which differential RNA expression downstream of Phf6 contributes to VEGF-A driven angiogenesis.

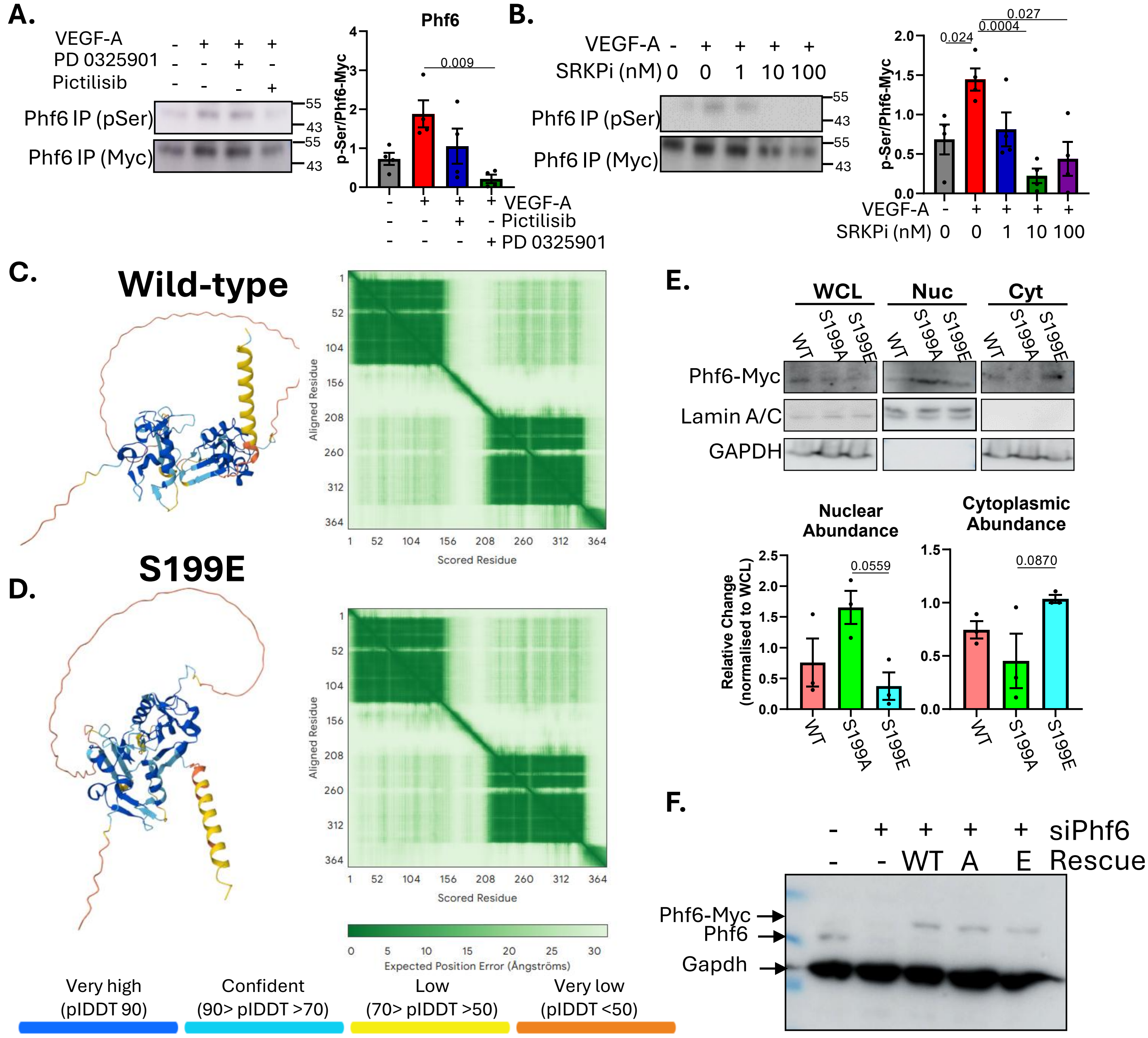


**Supplemental Figure 6.** A. Effects of AKT and ERK1/2 inhibition on Phf6 phosphorylation. Phf6 (Myc tagged) was overexpressed in MCECs in the presence of VEGF-A stimulation (24 h) and either PD325901 (Erk1/2 inhibitor) or Pictilisib (PI3K/AKT inhibitor) and immunoprecipitated from whole cell lysate. Western blots were used to detect the relative amounts of serine phosphorylation (n=4), significance assessed by one-way ANOVA, significance indicated. B. Effect of SRPK inhibition on Phf6 phosphorylation. Phf6 (Myc tagged) was overexpressed in MCECs in the presence of VEGF-A stimulation (24 h) and increased concentrations of the SRPK inhibitor MSC1186 (SRPKi). and immunoprecipitated from whole cell lysate. Western blots were used to detect the relative amounts of serine phosphorylation (n=4), significance assessed by one-way ANOVA, significance indicated. A-B. Predicted Phf6 structures for wild type (A) and phospho-mimetic (S199E) Phf6 (B), generated by alphafold, along with alphafold generated scoring of predicted residue alignment. C. Wild-type, phospho-deficient (S199A) and phospho-mimetic (S199E) Phf6 were overexpressed in MCECs for 48 hours followed by nuclear-cytoplasmic fraction and Western blot analysis (GAPDH and Lamin A/C were included as cytoplasmic/nuclear markers respectively). n=3, significance assessed by one-way ANOVA, significance indicated. F. Validation of Phf6 knock-down and rescue. Phf6 was knocked down with siRNA and rescued with either wild type (WT), S199 phospho-deficient (A) or S199 phosphomimentic (E) constructs (all Myc tagged) followed by Western blot to assess knock-down and rescue efficiency.

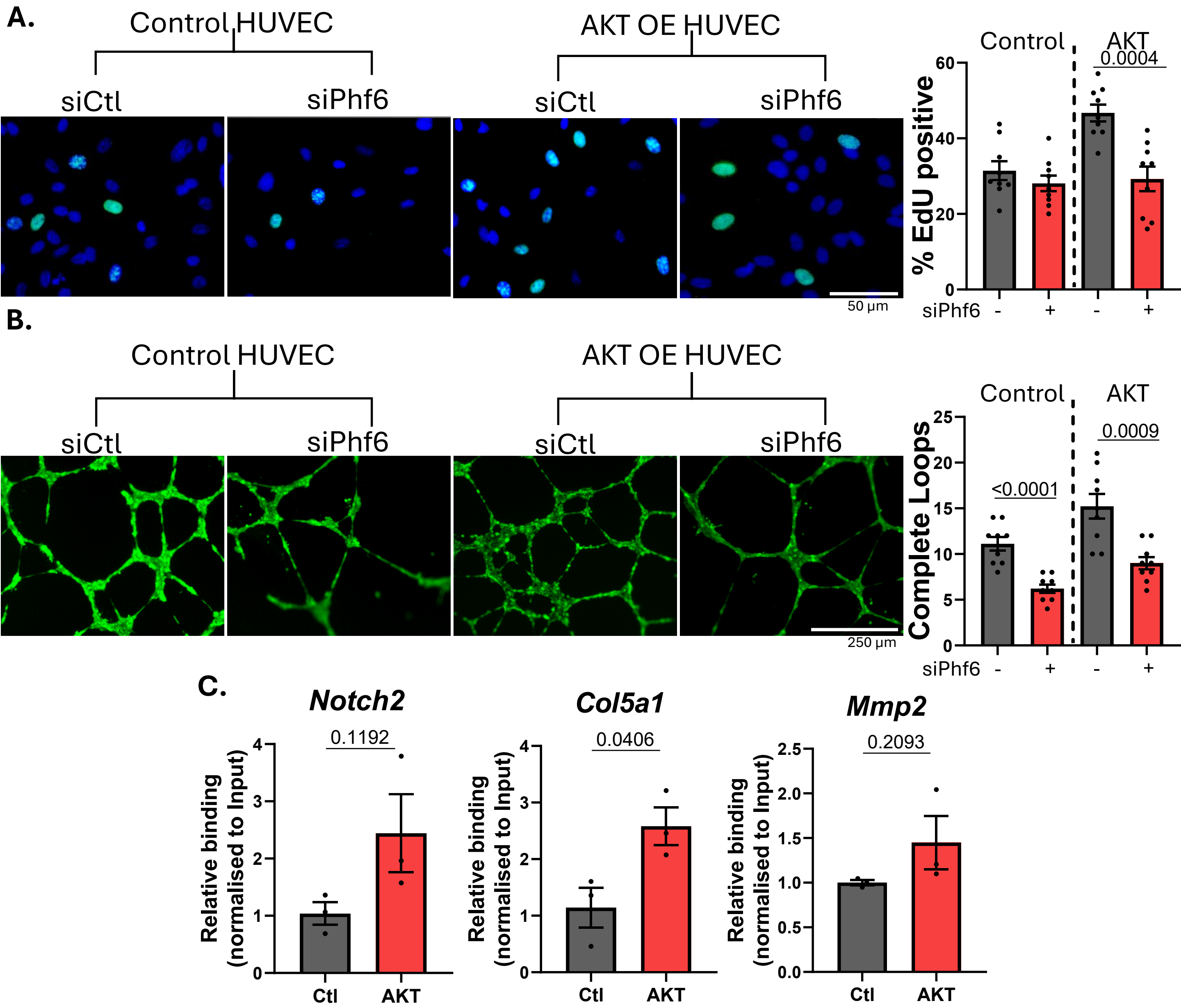


**Supplemental Figure 7.The effects of Phf6 knock-down in HUVECs overexpressing AKT.** A. Phf6 was knocked-down (siPhf6, 48 hours) in HUVECs constitutivley overexpressing AKT (vs. Wild-type controls) followed by an EdU incorporation assay (EdU green, DAPI blue) to assess the relative proliferation. Quantifications represent the % EdU positive cells. N=3 triplicates, quantifications show average ±SEM, significance assessed by one-way ANOVA, significance indicated. B. Phf6 was knocked-down (siPhf6, 48 hours) in HUVECs constitutivley overexpressing AKT (vs. Wild-type controls) followed by a tube formation assay. Quantifications represent the number of complete loops. N=3 triplicates, quantifications show average ±SEM, significance assessed by one-way ANOVA, significance indicated. C. Phf6 was immunoprecipitated from HUVECs constitutivley overexpressing AKT (vs. Wild-type controls) followed by RNA immunoprecipitation and qPCR analysis of the relative binding to selected targets of Phf6. Binding normalised to expression in matched inputs. N=3, quantifications show average ±SEM, significance assessed by students t-test, significance indicated.

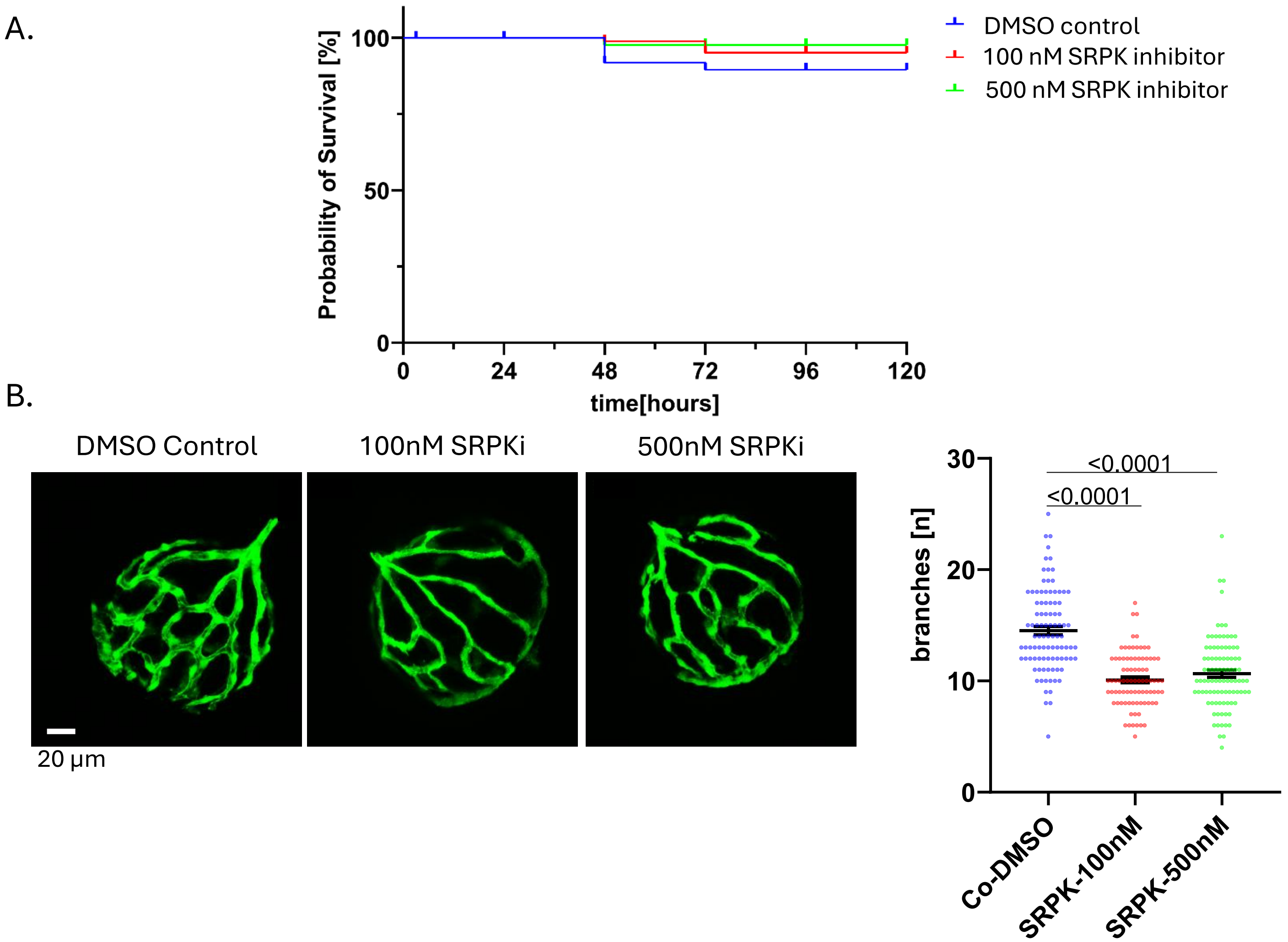


**Supplemental Figure 8. The effects of SRPK inhibition in zebrafish.** A. Mortaility of zebrafish embryos up to 120 hpf with incubation with either 100 nM or 500 nM of the SRPK inhibitor MSC-1186 in the culture water and B. assessment of the hyaloid vasculture formation (Fli1EGFP) following the incubation. N= DMSO Control: 95, 100nM SRPK inhibitor: 85 and 500nM SRPK: 94.

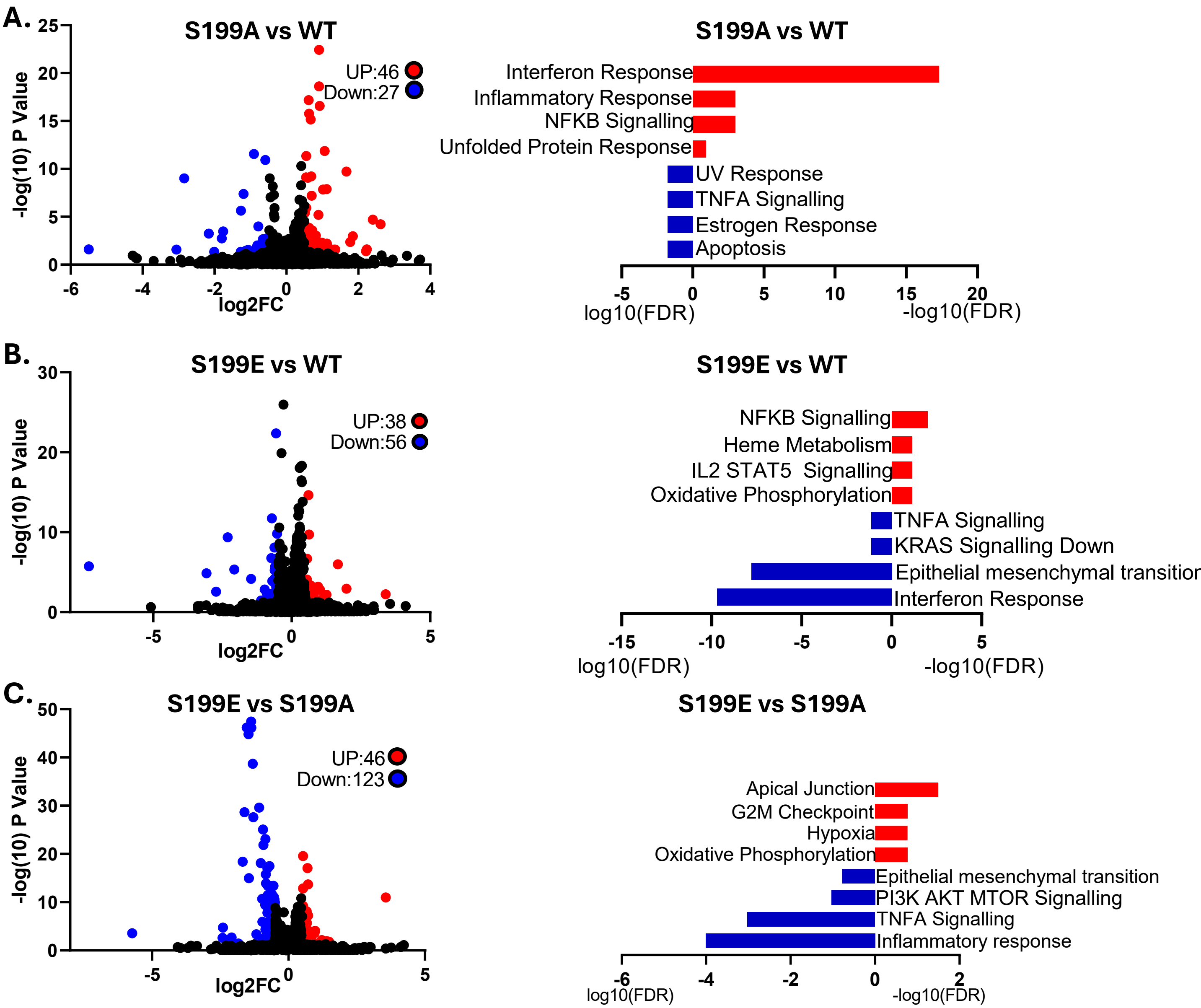


**Supplemental Figure 9. Extended analysis of the effects of Phf6 phospho-mutant overexpression on RNA expression.** A. Volcano plot of differentially expressed RNAs (Log2 FC >0.5, p. value <0.05), and the enriched MSigDB terms (red for up- and blue for downregulated in S199A vs. WT), among differentially expressed RNAs, following overexpression of phosphodeficient Phf6 (S199A) compared to wild-type (WT). B. Volcano plot of differentially expressed RNAs (Log2 FC >0.5, p. value <0.05), and the enriched GO biological process terms among differentially expressed RNAs, following overexpression of phosphomimetic Phf6 (S199E) compared to wild type (red for up- and blue for downregulated in S199E vs. WT). C. Volcano plot of differentially expressed RNAs (Log2 FC >0.5, p. value <0.05), and the enriched GO biological process terms among differentially expressed RNAs, following overexpression of phosphomimetic Phf6 (S199E) compared to overexpression of phosphodeficient Phf6 (S199A) (red for up- and blue for downregulated in S199E vs. WT).

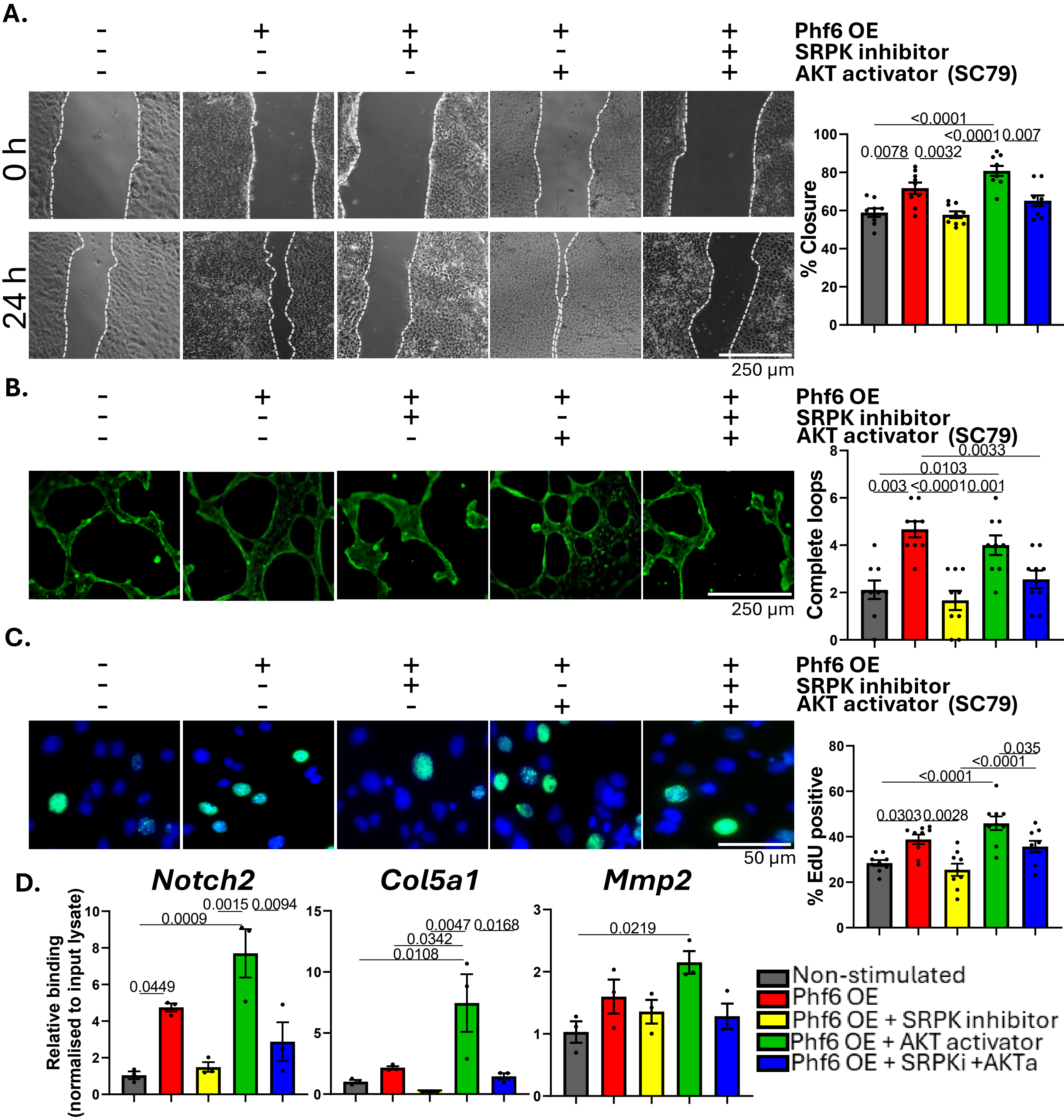


**Supplemental Figure 10. The role of SRPK kinase as an upstream regulator of Phf6.** A,B,C. Scratch migration assay (A), tubule formation assay (B) and EdU incorporation/proliferation assay (C) and quantifications, after Phf6 overexpression (48 hours) in the presence of SC79 (AKT activator,10 μM, 24 hours) AKT and MSC1186 (SRPKi, 10 nM 24 hours) as an inhibitor of SRPK. Each was done three times in triplicate, quantifications show average ±SEM, significance assessed by one-way ANOVA, significance indicated. D. RIP-qPCR analysis of the relative binding of Phf6 to selected targets after Phf6 overexpression (48 hours) in the presence of SC79 (AKT activator,10 μM, 24 hours) AKT and MSC1186 (SRPKi, 10 nM 24 hours) as an inhibitor of SRPK. N=3, quantifications show average ±SEM, significance assessed by one-way ANOVA, significance indicated.

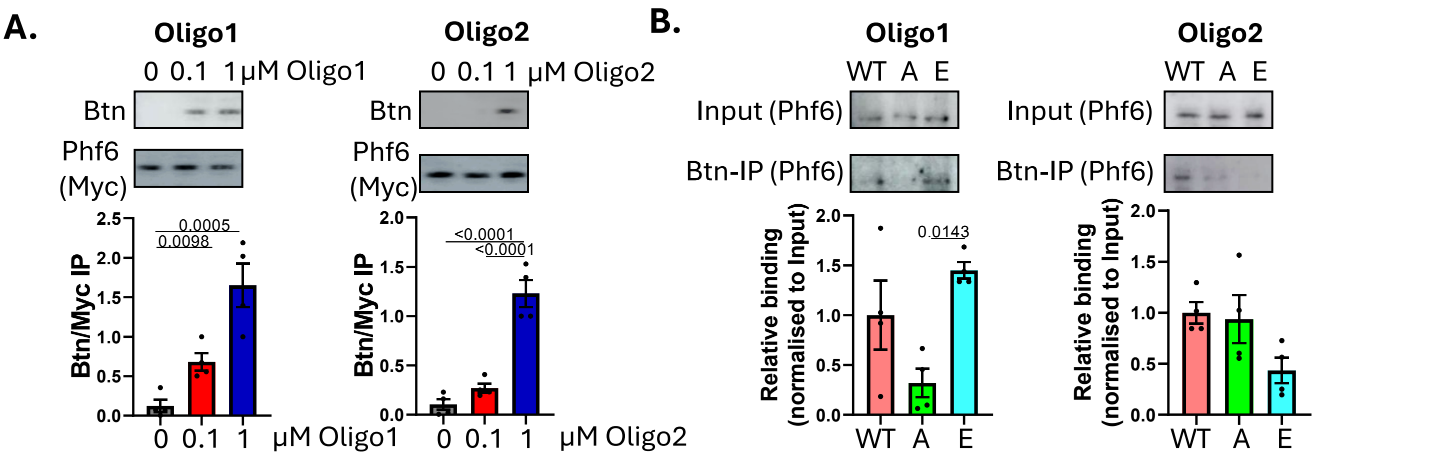


**Supplemental Figure 11. Optimisation and characterisation of competitive oligonucleotide inhibitors of Phf6 (extended from Figure 7 D-F).** A and B. Relative interaction of Phf6 with Oligo1 or Oligo2. A. Cells were transfected with wild-type (Myc-tagged, 48 hours) and either Oligo1 or Oligo2 (24 hours), followed by UV crosslinking, Phf6-Myc IP and Western blot. N=4, quantifications show average ±SEM, significance assessed by one-way ANOVA, significance indicated. B. Cells were transfected with wild-type (Myc-tagged, 48 hours) and either Oligo1 or Oligo2 (24 hours, 1 µM), followed by UV crosslinking, streptavidin capture and Western blot. N=4, quantifications show average ±SEM, significance assessed by one-way ANOVA, significance indicated.
